# Major effector loss reveals compensatory pathogenicity networks in a necrotrophic wheat pathogen

**DOI:** 10.64898/2026.09.14.751338

**Authors:** Shota Morikawa, Callum J. Verdonk, Leon V. Lenzo, Huyen T. T. Phan, Eiko Furuki, Kristina K. Gagalova, Johannes W. Debler, Chala J. Turo, Carl J. Mousley, Evan J. John, Bernadette M. Henares, Kar-Chun Tan

**Author notes:** These authors contributed equally (Shota Morikawa, Callum J. Verdonk).

## Abstract

Necrotrophic effectors (NEs) are key determinants of virulence in the necrotrophic fungal pathogen *Parastagonospora nodorum* that causes septoria nodorum blotch of wheat. However, targeted removal of three important NEs SnToxA, SnTox1, and SnTox3 in the mutant *Δtoxa13* previously revealed a redundancy mechanism is triggered, whereby pathogenicity on wheat is maintained. In this study, we investigated the gene regulatory profile underpinning this phenomenon and discover that virulence is not dependent on a fixed set of dominant effectors but instead arises from a flexible, epistatic compensatory network. Although host transcriptional responses to the *P. nodorum* wildtype SN15 and *Δtoxa13* infection remained largely conserved, consistent with an overlapping disease-susceptibility pathway, a significant upregulation of candidate effector genes was observed in *Δtoxa13*. This included the recently characterised NE *SnTox267*, and several other candidate effectors able to induce necrosis in the non-host *Nicotiana benthamiana*, each carrying a predicted structural fold conserved across other pathogens. We therefore provide further direct evidence that virulence is maintained in *P. nodorum* lacking three NEs by an epistatic and compensatory effector network, underpinned by changes in pathogen gene expression. Targeting conserved effector-mediated virulence mechanisms rather than individual host-specific gene-for-gene interactions may provide a more tractable route to host resistance.

## INTRODUCTION

Plant pathogenic fungi are broadly classified into three lifestyles: biotrophs, necrotrophs, and hemibiotrophs. Biotrophic pathogens derive nutrients from living host cells while minimising host detection, necrotrophs induce host cell death to access nutrients from dead tissue, and hemibiotrophs initially establish a biotrophic interaction before transitioning to a necrotrophic phase. Irrespective of lifestyle, plant pathogens secrete effector molecules that manipulate host cellular processes to promote infection (Lo Presti et al., 2015, Jaswal et al., 2020, McDonald et al., 2023). In necrotrophic fungi, proteinaceous necrotrophic effectors (NEs) can directly or indirectly interact with host susceptibility/sensitivity genes, triggering programmed cell death that favours, rather than restricts, pathogen proliferation (Tan et al., 2010, Kanyuka et al., 2022, Derbyshire and Raffaele, 2023b). Necrotrophic effector discovery has had a significant agricultural impact, as the identification of effector-host interactions has enabled the development of disease control strategies through the removal of host susceptibility factors in breeding programmes to improve the host resistance (McDonald and Solomon, 2018, Vleeshouwers and Oliver, 2014, McDonald, 2025).

*Parastagonospora nodorum* is a necrotrophic fungal pathogen and the causal agent of septoria nodorum blotch (SNB) of wheat (*Triticum aestivum*), a disease capable of causing yield losses of up to 30% (Murray and Brennan, 2009). Despite the deployment of control measures, SNB remains prevalent in wheat-growing regions worldwide (Lenzo et al., 2026). To date, several NEs have been characterised in *P. nodorum*, including SnToxA, SnTox1, SnTox267, SnTox3, and SnTox5. Deletion of individual NE genes typically results in reduced virulence on wheat cultivars carrying the corresponding susceptibility (S)-genes (Kariyawasam et al., 2023, Friesen et al., 2008b, Friesen et al., 2008a). Nevertheless, although NE-host interactions are qualitatively defined at the molecular level, disease development in the *P. nodorum*-wheat pathosystem is quantitatively inherited. Moreover, compensatory mechanisms among NE genes have been reported in multiple *P. nodorum* strains, which we have termed epistasis; whereby deletion or reduced expression of one effector can lead to increased expression of others (Tan et al., 2015, Phan et al., 2016, Richards et al., 2022). This phenomenon has also been observed for effector-host gene interactions in other pathosystems such as *Leptosphaeria maculans*-canola (Plissonneau et al., 2016) and *Pyrenophora tritici-repentis*-wheat (Manning and Ciuffetti, 2015). However, the molecular mechanisms triggering the changes in effector expression pertaining to epistasis remain undiscovered.

The Australian *P. nodorum* reference strain SN15 carries SnToxA, *SnTox1*, *SnTox3* and *SnTox267*, all of which are highly expressed during the early stage of plant infection, coinciding with host penetration (Ipcho et al., 2012, Rybak et al., 2017). Genes encoding these three NEs are also highly prevalent within the Australian *P. nodorum* population, present in over 96% of all isolates sampled (McDonald et al., 2013). Although NEs are primary determinants of disease in *P. nodorum*, they appear to function within an epistatic network – whereby elevated gene expression of some NEs compensates the loss of another in the event of NE gene loss or changes in host S-gene profile (John et al., 2022, Phan et al., 2016). In the *P. nodorum* SnTox1 deletion mutant toxa1-6, whereby the SnTox1-Snn1 susceptibility interaction is abolished, SnTox3 contributes to more disease at the Tox3-dependent SNB quantitative trait loci (QTL) QSnb.fcu–5BS (Snn3D1) (Phan et al., 2016). Surprisingly, simultaneous dual deletion mutants of SnToxA/SnTox3 or SnTox1/SnTox3 result in little to no reduction in virulence (Tan et al., 2014). A triple deletion of SnToxA/SnTox1/SnTox3 (toxa13-6, henceforth: Δtoxa13) does not reduce virulence on a collection of commercial wheat cultivars through conventional SNB infection, but virulence is increased on some lines (Phan et al., 2018). Neither did the necrosis-inducing activity of fungal culture filtrate significantly reduce in Δtoxa13 relative to the wildtype SN15 (Tan et al., 2015). Infections with Δtoxa13 have also been associated with the detection of minor SNB QTL (Phan et al., 2018), independent of other NEs. Together, these observations suggest that *P. nodorum* activates and relies on compensatory pathogenicity mechanisms to maintain virulence when major NEs are absent.

In this study, we investigate the epistatic nature of pathogenicity factors in the *P. nodorum* SNB pathosystem. Utilising comparative RNA sequencing (RNA-seq), we show that the Δtoxa13 mutant causes SNB on wheat by inducing the same conserved host-specific biochemical pathways as compared to the wildtype SN15. On the other hand, 60 candidate effector genes including the recently characterised NE SnTox267 (Richards et al., 2022) were significantly upregulated in the Δtoxa13 mutant relative to SN15. SnTox267 transcriptional upregulation correlated with increased disease contribution against a SnTox267-segregating wheat population, and 51% of tested commercial cultivars were sensitive to infiltrated SnTox267. Functional characterisation of a panel of the remaining effector candidates associated with this compensatory response revealed no activity when heterologously infiltrated into wheat directly but instead showed non-host necrosis-inducing capabilities in the model *Nicotiana benthamiana*, thus highlighting another layer of genetic control of host specificity associated with *P. nodorum*. Our results define a suite of pathogenicity-associated and non-host effector candidates that are upregulated in *P. nodorum* Δtoxa13, explaining disease persistence in the absence of major NEs and supporting a flexible, epistatic effector network. This framework provides a basis for systematically mapping interactions within a compensatory effector-regulatory network, in which coordinated changes in effector expression collectively sustain virulence across pathosystems, with implications for prioritising the removal of dominant S-genes in crop breeding.

## MATERIAL AND METHODS

### Strains and cultures

*P. nodorum* strains used in this study are listed in **Table 1**.

**Table 1:** *P. nodorum* strains used in this study.

| Strain | Description | Source |
| --- | --- | --- |
| SN15 | *P. nodorum* wildtype | Department of Primary Industries and Regional Development, Western Australia |
| *toxa-18* | SN15 with a deletion of *SnToxA* | (Friesen et al., 2006) |
| *tox1-6* | SN15 with a deletion of *SnTox1* | (Phan et al., 2016) |
| *tox3-30* | SN15 with a deletion of *SnTox3* | (Tan et al., 2014) |
| *toxa1-3* | *toxa-18* with a deletion of *SnTox1*;<br>*SnToxA/SnTox1* double-deletion | This study |
| *toxa3-10* | *toxa-18* with a deletion of *SnTox3*;<br>*SnToxA/SnTox3* double-deletion | (Tan et al., 2015) |
| *tox13-22* | *tox3-30* with a deletion of *SnTox1*;<br>*SnTox1/SnTox3* double-deletion | (Tan et al., 2014) |
| $\Delta toxa13$ | Original name *toxa13-6*: *toxa3-10* with a deletion of *SnTox1*;<br>*SnToxA/SnTox1/SnTox3* triple-deletion. | (Tan et al., 2015) |
| $\Delta tox267$ | SN15 with a deletion of *SnTox267* | This study |

### Generation of *P. nodorum* effector gene knockout mutants

Knockout constructs of SnTox1 or SnTox267 were constructed via fusion PCR (Hilgarth and Lanigan, 2020), and the *P. nodorum* toxa-18 (for SnTox1 deletion in toxa1-3) or SN15 (for SnTox267 deletion in Δtox267) was transformed as described previously (Morikawa et al., 2026).

### RNA extraction, sequencing and quality control of in-vitro and in-planta samples

RNA-seq data were generated previously using methods described in (Jones et al., 2019), using three-day post-infected lesions excised from detached wheat leaves (Rybak et al., 2017). RNA from SN15 (n = 4) and toxa13-6 (Δtoxa13) (n = 3) grown under in-vitro and in-planta (cv. Halberd; Tsn1/Snn1/Snn3) conditions were processed as detailed in (Jones et al., 2019). Tween20-treated (n = 3) leaves were used as uninfected control for wheat transcriptome. Library preparation and sequencing were performed by the Ramaciotti Centre of Genomics (The University of New South Wales).

### Differential gene expression analysis

For host analysis, raw in-planta reads were processed as described at https://github.com/LeonLenzo/a13. Briefly raw reads were passed through BBrepair (https://sourceforge.net/projects/bbmap/) to remove broken pairs, fastp (Chen et al., 2018) to remove low quality bases, kallisto (Bray et al., 2016) for pseudoalignment to Chinese Spring (IWGSC 2.1 Ref Seq, GCF_018294505.1) cDNA (Zhu et al., 2021), DESeq2 v1.48.2 (Love et al., 2014) for differential gene expression analysis, using a stringent cutoff for absolute LFC of ≥2 (T. aestivum) (Krattinger et al., 2019) and a Benjamini-Hochberg adjusted p-value of <0.05. The package clusterProfiler (v4.16.0) (Wu et al., 2021) was used for gene ontology (GO) enrichment with annotations pulled from Ensembl BiomaRt (v2.1), before manual curation into biologically relevant subcategories for visualization using ggplot2 (Wickham, 2016).

For pathogen analysis, FastQC v0.11.5 (http://www.bioinformatics.babraham.ac.uk/projects/fastqc) was utilised for quality control and Trimmomatic v0.39 (Bolger et al., 2014) was used for trimming the sequencing adapters. In-planta and in-vitro reads were filtered using BBsplit from the BBmap suite v39.84 (https://sourceforge.net/projects/bbmap/) with the *P. nodorum* SN15 reference genome (Bertazzoni et al., 2021) to retain RNA sequences belonging to the fungus. Filtered reads were aligned using STAR v2.6.4 (Dobin et al., 2013) and mapped to the *P. nodorum* SN15 genome. Aligned reads were quantified using the SubRead featureCount v2.0.0 package (Liao et al., 2014). Genes with more than five mapped reads (*P. nodorum* SN15) were retained. Gene ontology (GO) for the SN15 genome was assigned through InterProScan 5 (Jones et al., 2014, Blum et al., 2025). Fungal gene differential expression was determined by the R package DESeq2 v1.36.0 (Love et al., 2014). The noise of the Log_2_ fold change (LFC) of genes was reduced using the R package apeglm called through DESeq2 (Zhu et al., 2019). The cutoff for differential expression was an absolute LFC of >1 (*P. nodorum*) with a Benjamini-Hochberg adjusted p-value of <0.05. GO terms overrepresented in Δtoxa13 were determined using GOseq v1.26.0 (Young et al., 2010). Differential expression and GO terms can be found in **Supplemental File S1**. The sixty identified *P. nodorum* candidate effector genes upregulated in the in-planta RNA-seq analysis (LFC > 1; adjusted p < 0.001) were defined by the product containing signal peptide sequences predicted using SignalP 6.0 (Teufel et al., 2022) and being >80 amino acids (AA) and <350 AA in size. *P. nodorum* genes were categorised into down-regulated, upregulated and unchanged in-planta of Δtoxa13 and the upstream regions containing the promoters of the categorised genes were extracted using Bedtools (Quinlan and Hall, 2010).

### Infiltration of proteins & candidate lysates

SnTox267 was expressed, purified and infiltrated into wheat leaves as previously described (Phan et al., 2026). Candidate effectors culture filtrates were expressed in Pichia pastoris (syn. Komagataella phaffii) using the pGAPzα expression system as previously described (Phan et al., 2018). Culture filtrates were infiltrated into 12-day old wheat seedlings as previously described (Tan et al., 2015). Briefly, a one CC needleless plastic syringe was used to infiltrate each candidate effector culture filtrate suspension into the first leaf, with the extent of the leaf infiltration region marked with a non-toxic pen as previously described (Tan et al., 2014). SnTox3 was used as a positive control. Seven days following infiltration, the plants were visually evaluated for effector sensitivity (Tan et al., 2012). An empty-vector control was also used to establish a symptom baseline to evaluate possible damage due to the infiltration process.

### In-silico analysis of *P. nodorum* candidate effectors

AA sequences of *P. nodorum* candidate effectors were analysed on EffectorP 3.0 (Sperschneider and Dodds, 2022). Predector v1.2.7 (Jones et al., 2021) scores of *P. nodorum* candidate effectors were retrieved from (Jones et al., 2024). PHI-base was used to identify any characterised virulence-associated orthologues (Urban et al., 2025). Protein structures of SnTox1 and upregulated *P. nodorum* candidate effectors were predicted using AlphaFold 3 (Abramson et al., 2024). Signal peptides predicted using SignalP 6.0 were removed before the structural predictions (Teufel et al., 2022). Protein structures were visualised and analysed on Open-Source PyMOL v3.1 (https://github.com/schrodinger/pymol-open-source), and the predicted structures were coloured using pymol-color-alphafold (https://github.com/cbalbin-bio/pymol-color-alphafold).

The following analysis was performed on Geneious Prime v2024.0.4 (https://www.geneious.com) for GH11 xylanases. A custom BLAST database containing genomes from 17 fungal phytopathogens and four non-pathogenic fungi was created (**Supplemental Table S1**). PnXyn1 amino acid sequence was used as a query for a TBLASTN search (Camacho et al., 2009) using default parameters. Identified AA sequences of fungal PnXyn1 homologs, characterised fungal GH11 (glucoside hydrolase 11) xylanases, three characterised bacterial GH11 xylanases and the GH10 (glucoside hydrolase 10) xylanase ppxyn1 from the oomycete Phytophthora parasitica were aligned using the Clustal Omega algorithm (Sievers et al., 2011). A neighbour-joining distance tree (Saitou and Nei, 1987) was created with 1000 bootstraps and ppxyn1 as the outgroup.

### Agroinfiltration of candidate effectors of *P. nodorum*

Agroinfiltration of *P. nodorum* candidate effectors was performed as previously described with some modifications (Debler et al., 2021, Debler et al., 2026). Nicotiana benthamiana was grown in potting soil (RichGro) at 18 – 25 °C under a 12 hr: 12 hr light: dark regime for approximately 6 weeks before agroinfiltration. Candidate effectors identified in *P. nodorum* without the native signal peptide flanked by the pDONR attB1 and attB2 sequences were ordered individually as a synthetic DNA gBlocks (Integrated DNA Technologies) (**Supplemental Table S2**). The candidate effector genes were cloned into pDONRZeo, then into both pEAQ-HT-DEST1-His (for cytoplasmic expression) and pEAQ-HT-DEST1-PR1-His (carrying the Medicago truncatula PR-1 signal peptide to target the protein to the apoplast) via Gateway recombination using manufacturer’s instructions (Thermo Fisher Scientific). Agrobacterium tumefaciens strain AGL1 was transformed via electroporation using a MicroPulser Electroporator (Bio-Rad) using the preset protocol “Agr”. A. tumefaciens AGL1 transformants carrying the plasmids pEAQ-HT-DEST1-PR1-His containing either GFP and Ascochyta rabiei NLP2 (necrosis-and ethylene-inducing peptide (Nep1)-like protein 2) were used as a negative and positive control, respectively (Debler et al., 2021). A. tumefaciens transformants carrying the correct plasmids were grown in LB media supplemented with 30 µg/mL kanamycin and 50 µg/mL rifampicin at 200 rpm at 27 °C for approximately 40 hr. The A. tumefaciens cells were pelleted at 3000 g for 5 min, then resuspended in 10 mL Induction Buffer (10 mM MES, 10 mM MgCl_2_, pH 5.7), and the OD_600_ was measured using a BioTek Synergy HTX Multimode Reader (Agilent Technologies). The A. tumefaciens cells were pelleted again and then resuspended in Induction Buffer supplemented with 400 µM acetosyringone to an OD_600_ of approximately 1.0. The A. tumefaciens suspension was cultured at 150 rpm at 27 °C for 3 hr. The adaxial side of broad N. benthamiana leaves was infiltrated with the A. tumefaciens suspension using a 1.0 mL needleless syringe. N. benthamiana leaves were excised and photographed 10 days post-agroinfiltration. GFP fluorescence and necrosis were visualised on the ChemiDoc™ XRS+ System with Image Lab™ software (Bio-Rad) using excitation wavelengths between 460 – 490 nm and 520 – 545 nm and emission wavelengths of 530 nm and 605 nm, respectively.

### Association mapping and quantitative trait loci (QTL) analysis

The 105:ZIF14 × 56:ZWB11 double-haploid wheat population (DH105×56), consisting of 241 progeny, was previously developed where 105:ZIF14 and 56:ZWB11 were weakly/insensitive (score <2) and strongly sensitive to SnTox267, respectively (Phan et al., 2026). SnTox267 sensitivity was mapped to 6 loci in this DH population; with Snn7 (Richards et al., 2022) and five other QTL on chromosomes 2D (long arm), and 2A2, 2A3, 2B1, 5B, 7B1, respectively (Phan et al., 2026). In this study, the DH105×56 population was used to conduct QTL mapping of seedling SNB from SN15 and Δtoxa13 to dissect the contribution of SnTox267 to SNB. Briefly, a total of 3,403 markers including DArTseq and SNP markers were used to construct a genetic map for the DH105×56 population (Phan et al., 2026). For infection, a whole plant spray using *P. nodorum* pycnidiospores was carried out on two-week old seedlings in randomised biological triplicates and SNB severity was visually assessed and scored on a scale of one to nine (Phan et al., 2016). A score of one indicates no disease symptoms, whereas a score of nine indicates a fully necrotised plant (Phan et al., 2016, Phan et al., 2018). QTL mapping was undertaken using MultiQTL software, v.2.5 (MultiQTL Ltd, Institute of Evolution, Haifa University). The QTL mapping was based on a genetic linkage map previously built for the DH105×56 population using the Kosambi mapping function using MultiPoint v.3.2 software (MultiQTL Ltd, Institute of Evolution, Haifa University) from maximum recombination frequencies of 0.5. Seedling disease scores were taken from the average of biological replicates for interval mapping. QTL with a LOD score ≥3.0 were declared as significant as previously described (Phan et al., 2026).

## RESULTS

### Comparative RNA-seq reveals limited divergence in wheat host transcriptome response to SN15 and Δtoxa13 infection

The direct and indirect targets of the NEs SnToxA, SnTox1 and SnTox3 are well documented, and all lead to susceptible wheat programmed cell death (Winterberg et al., 2014, Kariyawasam et al., 2023, Veselova et al., 2021). However, *P. nodorum* lacking all three of these major NEs (Δtoxa13) retained wildtype virulence on most wheat cultivars (Tan et al., 2015, Phan et al., 2018). Here, we attempted to uncover the potential redundancy products produced by Δtoxa13 for virulent factors. First, we wondered whether there was a difference in wheat host response during infection for SN15 and Δtoxa13. To investigate any potential difference in wheat host immune response, we performed a comparative RNA-sequencing approach on infected wheat seedlings to interrogate the differentially expressed (DE) genes between SN15 and Δtoxa13 infection, relative to uninfected leaves. A principal component (PC) analysis of the SN15 and Δtoxa13 host RNA-seq data revealed a clear segregation consistent across replicates (**Supplemental Figure S1**). Across both SN15 and Δtoxa13 infection (SN15 ∪ Δtoxa13), 19,381 (22%) of all reference wheat genes were DE during infection; and the majority (15.1%) of these total DE genes overlapped during SN15 and Δtoxa13 infection (SN15 X Δtoxa13) (**Figure 1A**). We interrogated the enrichment of gene ontology (GO) terms within the 1.8% subset of Δtoxa13 genes which were DE relative to SN15 (SN15’ X Δtoxa13) during infection, and found that relative to SN15, enriched Δtoxa13 GO terms do not appear to be independent of conventional effector-triggered susceptibility pathways leading to programmed cell death previously identified (Kariyawasam et al., 2023) (**Figure 1B**). Of note, the GO annotations associated with energy and photosynthesis; “photosystem I” and “photosystem II” were upregulated in Δtoxa13 compared with SN15, supporting the light-associated role of NEs within *P. nodorum* (Friesen and Faris, 2010, Richards et al., 2022, Breen et al., 2016, Gao et al., 2015). Given the marginal difference in host response between SN15 and Δtoxa13 infection on wheat, we suggest that Δtoxa13 carries conserved pathogenicity factors beyond SnToxA, SnTox1 and SnTox3 which utilise similar pathways as SN15 for eliciting host tissue death and subsequent infection.

**Figure 1:**
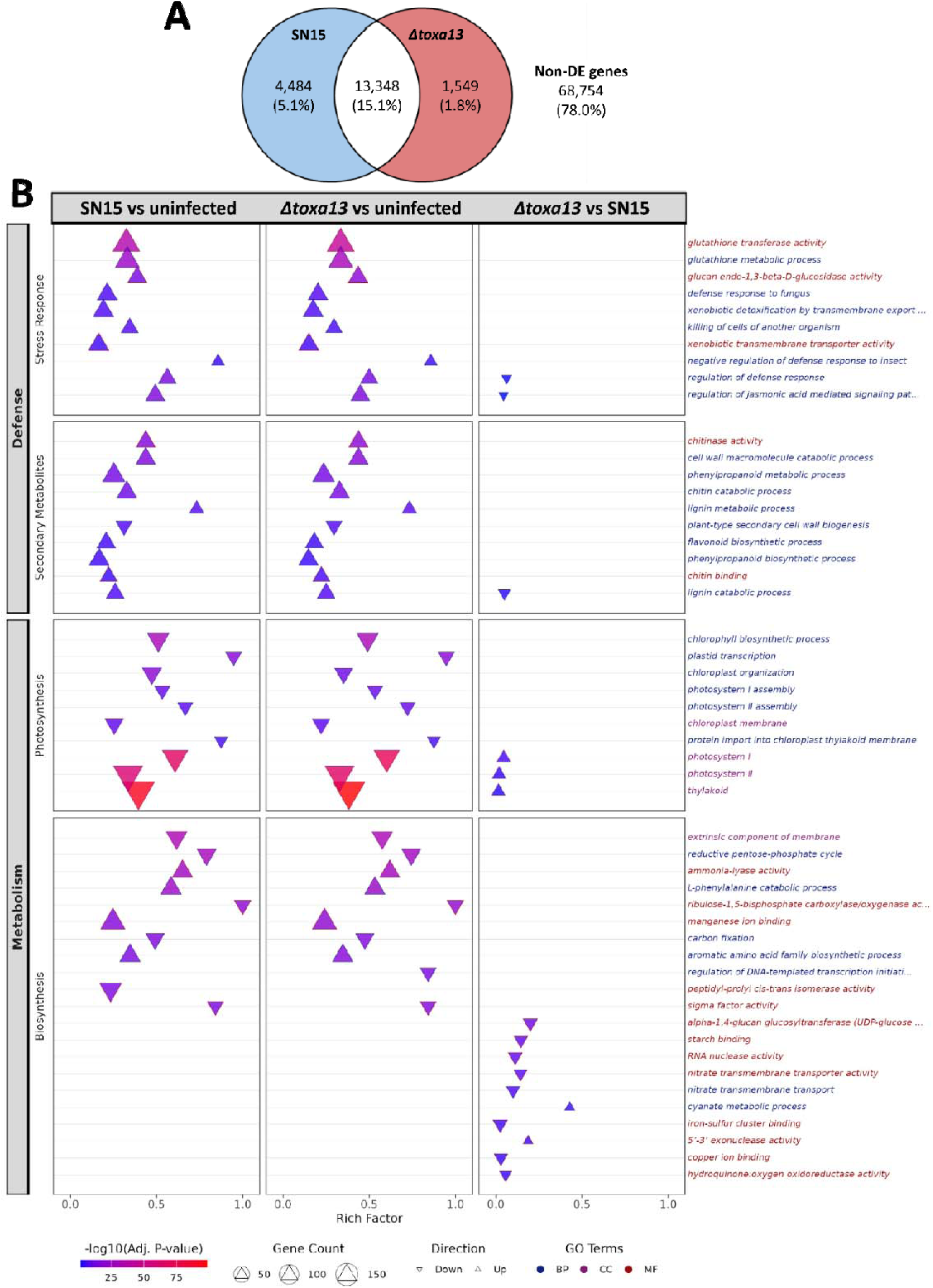
Comparative RNA-seq analysis of wheat transcriptome to SN15 and Δtoxa13 infection. **(A)** Differential analysis comparing total wheat genes (reference cv. Chinese Spring) during SN15 and *Δtoxa13* infection. **(B)** GO terms significantly enriched (FDR < 0.05) among wheat differentially expressed genes (|log2FC| ≥ 2) are shown for three pairwise comparisons: SN15 vs Uninfected (tween20), *Δtoxa13* vs Uninfected (tween20) and *Δtoxa13* vs SN15. GO terms were manually assigned into two categories (Defense and Metabolism), each with two subcategories (Stress Response/Secondary Metabolites and Photosynthesis/Biosynthesis) with top 10 significant terms per subcategory per comparison shown. Points are positioned on the x-axis by Rich Factor (ratio of differentially expressed genes to background genes in each GO term), with point size indicating gene count and point shape indicating direction of overall gene expression status (△ = upregulated, ⍰ = down-regulated). Point colour represents significance level (-log_10_ adjusted p-value). GO term labels on the right are color-coded by ontology: blue for Biological Process (BP), purple for Cellular Component (CC), and red for Molecular Function (MF).

### Candidate effector genes are upregulated in the absence of *ToxA*, *Tox1* and *Tox3*

As we could not observe distinct changes in GO representation profile of host-plant responses when comparing infection by SN15 and Δtoxa13, yet some NEs can be upregulated in the absence of others (Phan et al., 2016, Richards et al., 2022), we hypothesised that other upregulated genes in Δtoxa13 may encode virulence factors in *P. nodorum*. To identify *P. nodorum* genes encoding potential virulence factors, a comparative RNA-seq approach for both in-vitro and in-planta infection conditions was employed between SN15 and Δtoxa13. A PC analysis revealed the majority of expression variation (PC1: 93%) divides along in-vitro and in-planta growth conditions, but also that a distinct, albeit small fraction (PC2: 4%), clearly separated the expression profile of SN15 and Δtoxa13 (**Figure 2A**). This suggests marginal variance under in-vitro conditions, but revealing that host-associated conditions drive compensatory responses in Δtoxa13 to maintain virulence in the absence of major NEs. The in-vitro RNA-seq analysis revealed 742 down-regulated genes (4.5%) and 973 upregulated genes (5.9%) in Δtoxa13 compared to SN15. Meanwhile, the in-planta RNA-seq analysis revealed 441 down-regulated genes (2.7%) and 456 upregulated genes (2.8%) in Δtoxa13.

**Figure 2:**
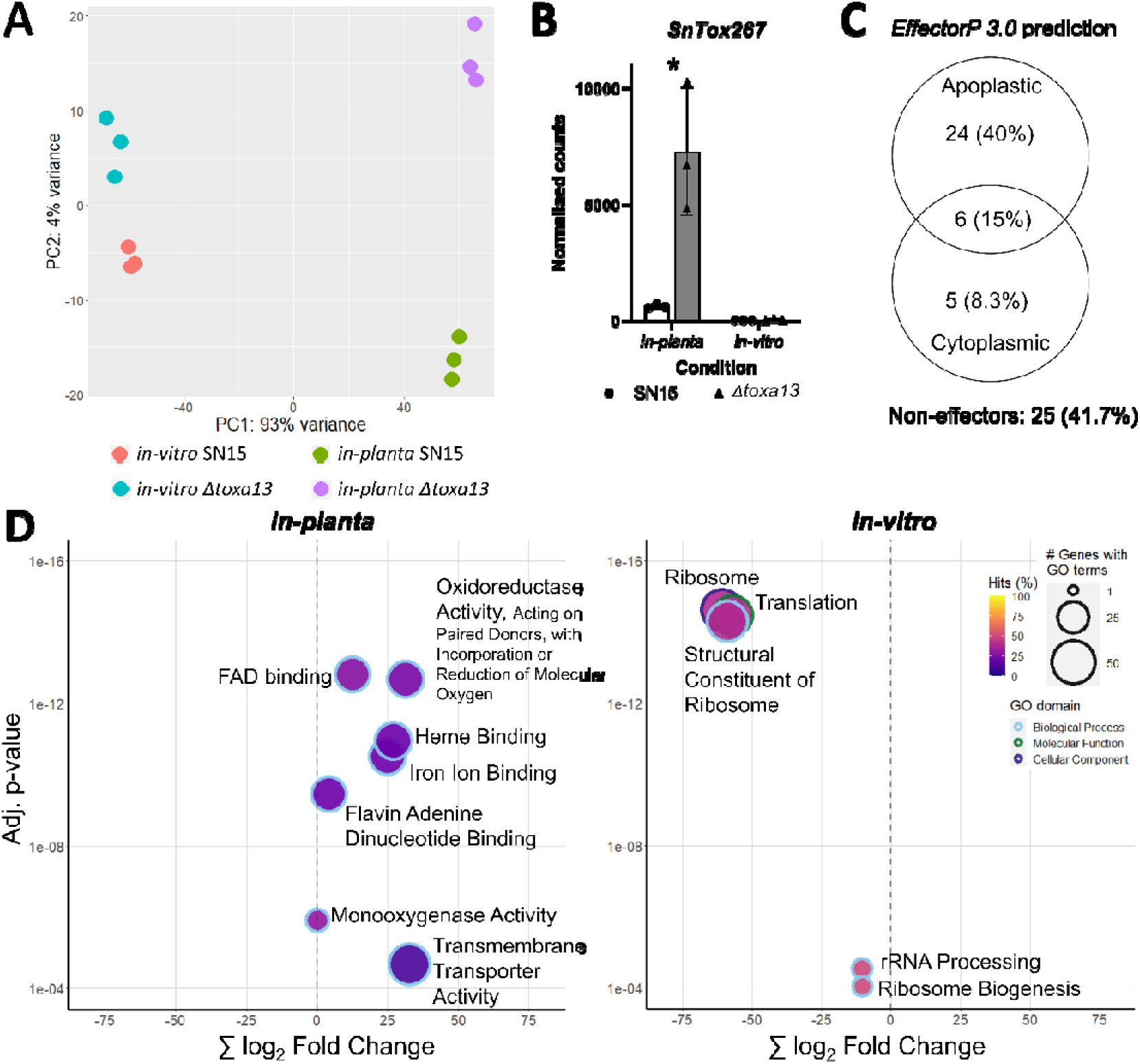
Comparative RNA-seq analysis between SN15 and Δtoxa13 during infection (in-planta) and during in-vitro conditions. **(A)** Principal component analysis (PCA) of transcriptomes of SN15 and *Δtoxa13*. **(B)** Transcriptome normalised counts for *SnTox267 in-vitro* and *in-planta* for SN15 and *Δtoxa13.* \**p-*value < 0.05. **(C)** Venn diagram for total *EffectorP 3.0* (Sperschneider and Dodds, 2022) prediction of all 60 upregulated candidate effectors identified in *Δtoxa13,* and their predicted localisation (apoplastic and/or cytoplasmic). **(D)** Gene-ontology (GO) enrichment analysis of differentially expressed genes in *Δtoxa13 in-planta* and *in-vitro* relative to SN15. There is an overall up-regulation of genes involved in catalysis *in-planta* while a down-regulation of genes related to translation *in-vitro*. The *x*-axis indicates the sum of the *Log_2_* fold change of differentially expressed genes in each GO term. The *y*-axis indicates the adjusted *p*-value.

Differential RNA-seq gene analysis indicated the NE SnTox267 was significantly upregulated in Δtoxa13 in-planta (**Figure 2B**). Including SnTox267, 60 genes encoding small putatively secreted proteins (PnSsp), which we have termed “candidate effectors”, were also transcriptionally upregulated in Δtoxa13 in-planta (**Supplemental Table 3**). This striking over-representation of secreted candidate effectors upregulated in Δtoxa13 in-planta (representing 13% of all upregulated genes) identifies them as the likely dominant response to effector loss and why we prioritised them in downstream analyses of compensatory virulence. Each of the candidate effectors was assessed in the effector prediction tool effectorP 3.0 (Sperschneider and Dodds, 2022), and 35/60 (58%) were predicted to harbour an effector function (**Figure 2C**). GO term enrichment revealed that Δtoxa13 displays distinct transcriptional responses relative to SN15 during in-planta infection versus in-vitro growth (**Figure 2D**). Based on a stringent set of candidate effector genes predicted previously (Jones et al., 2024), PnXyn1 and PnSsp25 were the two most upregulated candidates observed in Δtoxa13, and along with SnTox267, their up-regulation was exclusively observed in-planta and not in-vitro.

Interrogation of our 60-candidate effector list revealed the majority (47/60: 78%) were 100% present across a panel of *P. nodorum* 173 isolates (Jones et al., 2024). We wondered whether these candidate effector genes were maintained in *P. nodorum* as part of a conserved regulatory redundancy-based strategy, given upstream intergenic region (UIR) length is evolutionarily conserved in fungi and has been shown to strongly predict gene regulation and functional importance (Noble and Andrianopoulos, 2013). Analysis of all DE genes’ UIR (containing promoters) revealed that upregulated genes had significantly longer UIR than downregulated or non-differentially expressed genes, with median lengths of 865, 667 and 519 bp, respectively (**Supplemental Figure S2**). Among our candidate effector gene list, the median UIR length increased to 1296 bp.

### Epistatic up-regulation and contribution of SnTox267 to virulence in the absence of major effectors

As we observed a distinct epistasis for the remaining characterised NE in SN15, SnTox267, in our RNA-seq analysis (**Figure 2B**), we wondered whether we could validate the direct impact of effector epistasis on SnTox267 expression using a quantitative reverse-transcriptase PCR (qRT-PCR) assay. *P. nodorum* SN15 carrying selective deletions of either SnToxA (toxa-18), SnTox1 (tox1-6) or SnTox3 (tox3-30) showed upregulated levels of SnTox267 expression via qRT-PCR in-planta compared with wildtype SN15 (**Figure 3A**). Strains with dual-effector deletions: toxa1-3, toxa3-10 or tox13-22 showed further increases in SnTox267 expression relative to each single-effector deletion. Most interestingly, the triple NE deletion mutant Δtoxa13 had the highest relative SnTox267 expression, over six-fold higher relative to wildtype SN15. These data indicated that the observed upregulation of SnTox267 is not specific to individual NE loss but instead reflects a compensatory response that is progressively amplified with successive NE deletions.

**Figure 3:**
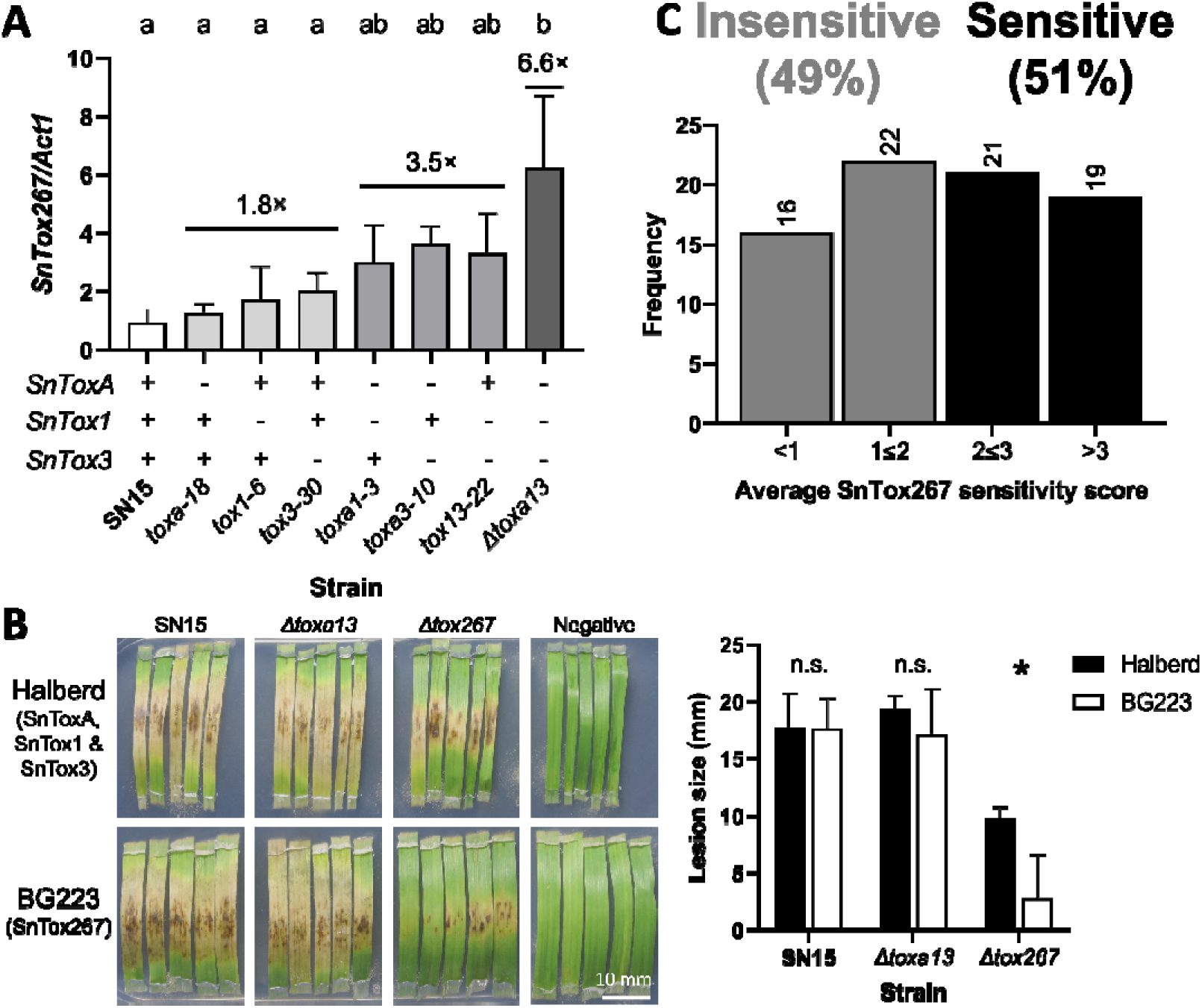
Deletion of SnToxA, SnTox1 and SnTox3 alleviate SnTox267 suppression. **(A)** Quantitative reverse-transcriptase PCR (qRT-PCR) assays show *SnTox267* is epistatic in single, double and triple-effector mutants in the *P. nodorum* SN15 background during wheat infection. Approximate mean fold-change relative to SN15 for each group shown above each bar. **(B)** Detached leaf assays indicate *Δtoxa13* can infect identically to SN15, while *Δtox267* shows reduced lesion sizes on the *SnTox267*-selective line BG223. Host-specific NE sensitivity shown in brackets under each corresponding wheat cultivar. \**p-*value < 0.05. **(C)** SnTox267 infiltrations into commercial cultivars suggest *SnTox267* sensitivity is present in half (51%) of all tested varieties. A score of 0 indicates insensitivity (no reaction); 1, slight chlorosis; 2, moderate chlorosis/slight necrosis; 3, moderate necrosis; 4, extensive necrosis. Accompanying scores for each wheat cultivar are available in **Supplemental Table S4**.

Given the upregulation of SnTox267 in NE-deleted variants and Δtoxa13, we decided to directly investigate the epistatic regulation of SnTox267. Firstly, the role of SnTox267-mediated host-response in Δtoxa13 infection was quantified relative to wildtype SN15. Detached leaf assays indicated that both SN15 and the triple-effector deletion Δtoxa13 caused comparable lesion sizes on the highly susceptible cv. Halberd (Tsn1, Snn1, Snn3), as well as on the SnTox267-selective line BG223 (tsn1, snn1, snn3) (**Figure 3B**). To compare, we generated a targeted SnTox267-deletion strain in the SN15 background. This SnTox267 knockout mutant, Δtox267, exhibited a 72% reduction in average lesion size on BG223 than on Halberd. The ability of SN15 and Δtoxa13 to successfully infect the wheat line BG223 demonstrates that SnTox267 is utilised during infection in both strains that was similarly observed in previous investigations (Phan et al., 2026, Phan et al., 2016). Interestingly, Δtox267 also had a 45 – 50% reduction in average lesion size on cv. Halberd compared to SN15 and Δtoxa13. This potentially suggests that SnTox267 may have a greater role in *P. nodorum* virulence than previously observed, at least when infecting the highly-susceptible wheat cv. Halberd.

To validate the function of SnTox267 and to phenotype cultivar sensitivity, we infiltrated SnTox267 into a collection of diverse commercial Australian wheat cultivars. We heterologously expressed and recombinantly purified SnTox267 (without its signal peptide) from Escherichia coli (Phan et al., 2026) and infiltrated into attached wheat leaves, scoring the severity of wheat host using a 0 (insensitive) to 4 (extensive necrosis) scale. Across the collection of 78 wheat cultivars, 51% were sensitive to recombinant SnTox267 with 19 (24%) showing strong chlorotic symptoms (average score >3) which indicated high susceptibility, while 38 (49%) cultivars were insensitive (average score <2) to SnTox267 infiltration (**Figure 3C** & **Supplemental Table S4**). Together, these results showed that SnTox267 elicits variable responses across commercial Australian wheat cultivars, with over half of the tested cultivars exhibiting susceptibility (sensitivity scores >2); at a much higher proportion than sensitivity to SnToxA or SnTox1 alone (Tan et al., 2014).

Next, we asked whether the epistatic up-regulation of SnTox267 increases it contribution to SNB disease phenotype. Therefore, interval mapping and SNB quantitative trait loci (QTL) analysis were undertaken using the markers previously developed for the double haploid 105ZIF14 × 56:ZWB11 (DH105×56) population, segregating for SnTox267 sensitivity (Phan et al., 2026), to explore any differences in the disease interactions produced by SnTox267 in SN15 and the Δtoxa13 mutant strain. Previous infiltration of the SnTox267 protein on the DH105×56 population revealed a major QTL on chromosome 2DL, where the S-gene Snn7 is located, explaining the highest phenotypic contribution at 18% in adult disease field trials (Phan et al., 2026). A QTL detected at chromosome 2D2 (Snn7) exhibited 13.8% of disease contribution during Δtoxa13 infection, higher than observed for SN15 at 10.3% (**Table 2**). We additionally observed a major QTL on 2A (15.6% of disease phenotype). This 2A QTL is similar to our previous observations with Δtoxa13 using a different marker set surrounding this region (Phan et al., 2016) and with previous epistatic studies on SnTox1 (John et al., 2022), along with responses to five diverse effector-containing *P. nodorum* isolates (at QTL 2AS) (Ara et al., 2026), suggesting further unidentified NEs may be contributing to SNB in parallel with the observed SnTox267-Snn7 interaction, or masked in the presence of SnToxA, SnTox1 or SnTox3 activity. Another minor QTL associated with Δtoxa13 only was observed on 6B, not observed for SnTox267 infiltration previously (Phan et al., 2026), although noted as an SNB QTL previously (Downie et al., 2020). It is likely that SnTox267 contributes marginally more to disease during Δtoxa13 infection than SN15, while several other minor QTLs emerge specific to Δtoxa13 infection.

**Table 2:**
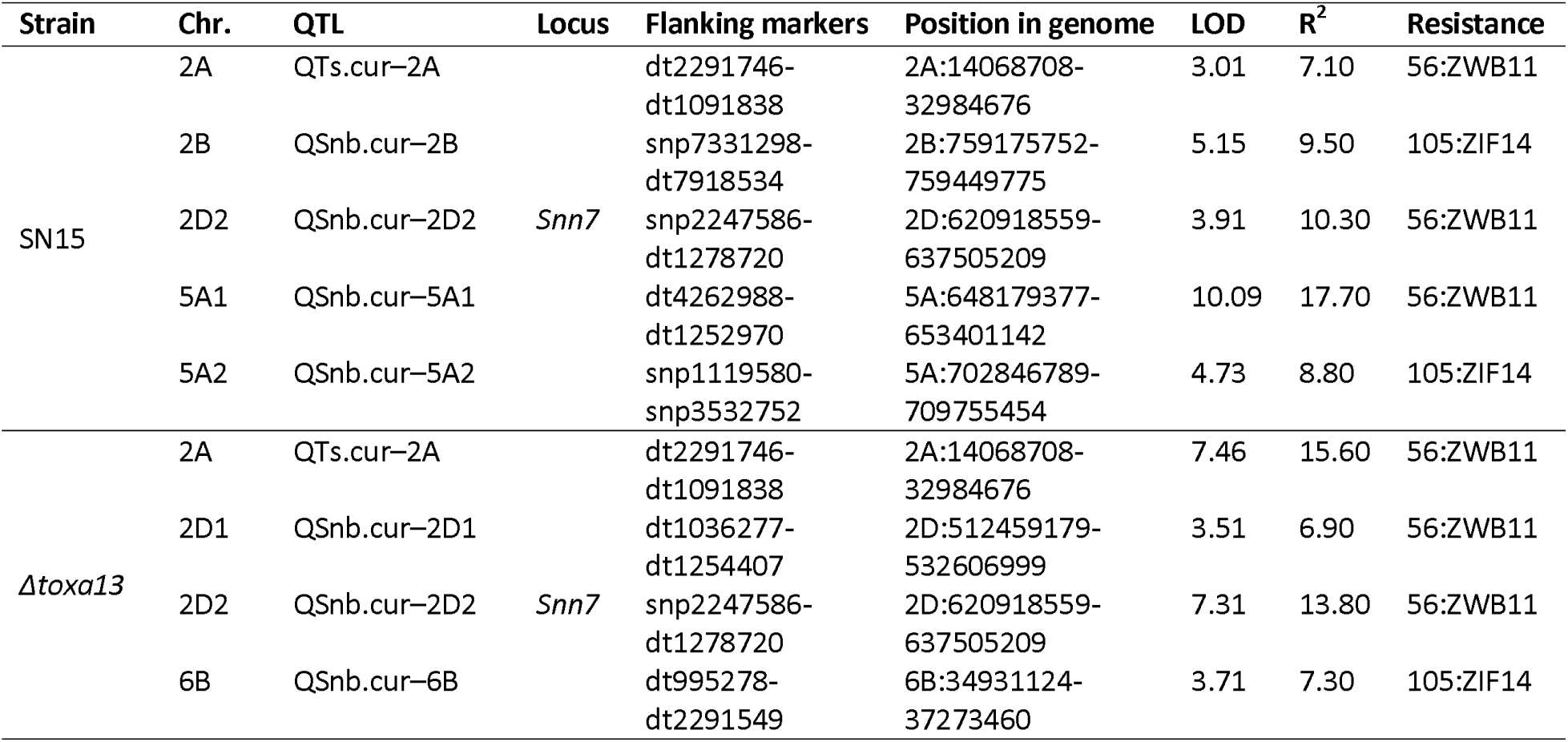
Summary of QTL detected from septoria nodorum blotch disease assessments at seedling growth stage using *P. nodorum* SN15 and Δtoxa13 on the DH105×56 population.

### Candidate effectors encoded by upregulated genes in Δtoxa13 are not host-specific but induce necrosis in N. benthamiana

Given the epistatic nature of NEs, we wondered whether our Δtoxa13-determined upregulated candidate effectors exhibited host-specific necrosis on wheat. To test host-specific activity for our candidate effectors, we expressed 14 of the top 17 candidates most highly expressed candidates from Δtoxa13 in Pichia pastoris. To observe any necrotic symptoms directly induced by each candidate effector, we initially infiltrated Pichia culture filtrates expressing each of the candidate effectors into four representative commercial wheat cultivars sensitive to SNB disease but with diverse genotypes and sensitivities to Δtoxa13 culture filtrates described previously (Tan et al., 2015): Halberd (Tsn1, Snn1, Snn3: sensitive to Δtoxa13 culture filtrate), Axe (Tsn1, Snn1, Snn3: insensitive to Δtoxa13 culture filtrate), Emu Rock (genotype predicted as tsn1, snn1, snn3: sensitive to Δtoxa13 culture filtrate), Mace (tsn1, snn1, Snn3: insensitive to Δtoxa13 culture filtrate). While we observed necrosis to SnTox3 as a positive control for all four wheat cultivars, we did not observe any host-specific necrosis for any of the other candidates tested, apart from PnSsp16, which induced necrosis on cv. Mace and Halberd (**Supplemental Figure S3**), although the development and appearance of symptoms were inconsistent.

As we were unable to consistently observe any host-specific necrosis activity for our candidate effector shortlist, we investigated whether these candidate effectors can elicit a non-host response (Kettles et al., 2017, Yang and Xu, 2026). Non-host activity can be assessed with Agrobacterium-mediated transient expression in the model dicot Nicotiana benthamiana (Yang and Xu, 2026), and we subsequently cloned our 14 wheat-assessed candidates into Agrobacterium. As *P. nodorum* utilises both apoplastic and cytoplasmic effectors (Kariyawasam et al., 2023), we investigate both localisations during N. benthamiana transient expression. Seven (50%) of the 14 candidates transiently expressed in N. benthamiana induced observable symptoms, five of which exhibited strong necrosis comparable to the NLP2 positive control (**Table 3**; **Figure 4; Supplemental Figure S4**). Of these seven positive necrosis-inducing hits, six acted in the cytoplasm – inducing necrosis in the absence of any localisation signal peptide. The only apoplast-only necrosis-inducing candidate was PnGlc1, which is predicted to be a glucanase, and may target conserved plant cell-wall components and potentially explain why it was unable to induce necrosis when localised in the cytoplasm. These data suggest that transcriptionally upregulated effector candidates in Δtoxa13 exhibited necrosis-inducing activity in N. benthamiana and may act as non-host effectors utilised in the absence of the primary NEs SnToxA, SnTox1 and SnTox3.

**Figure 4:**
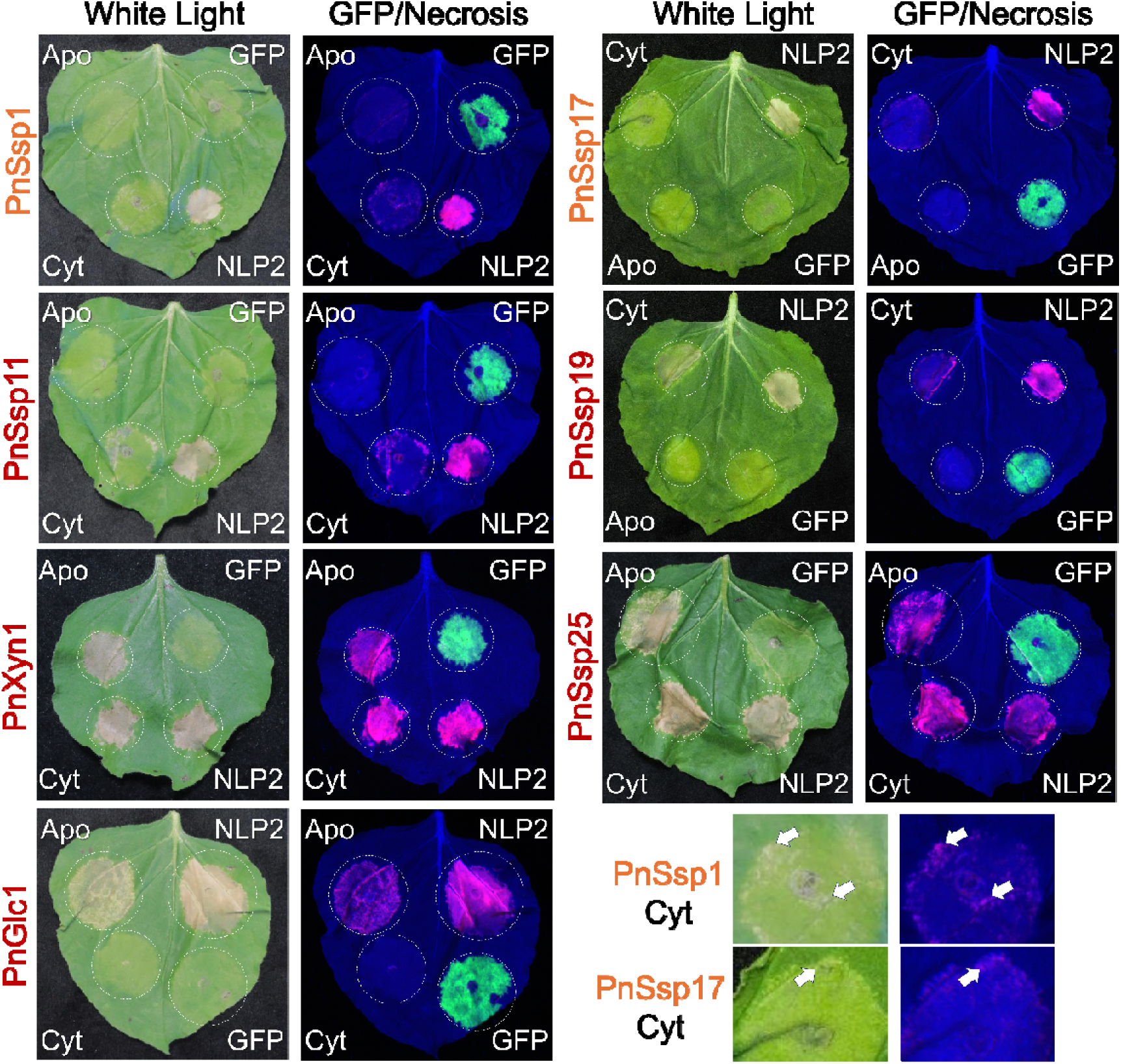
In-planta Δtoxa13 upregulated candidate effector genes induce necrosis on non-host Nicotiana benthamiana. Agroinfiltration of necrosis inducing up-regulated candidate effectors from *P. nodorum* Δtoxa13 and expressed in SN15. Candidate effectors with strong (red) and weak (orange) necrosis-inducing activity shown. Other non necrosis-inducing candidate effectors are available in **Supplemental Figure S4**. “White light” shows representative N. benthamiana leaves agroinfiltrated with one of 14 candidate effectors and photographed 10 days post-infiltration. NLP2 (necrosis-and ethylene-inducing peptide 1 (NEP1)-like protein 2) and GFP (green fluorescent protein) are the positive and negative controls, respectively. The location of each agroinfiltration was partially randomised. “Necrosis/GFP” shows the same leaf highlighting necrosis (pink) and GFP fluorescence (green). Cytoplasmic (“Cyt”) and apoplastic (“Apo”) localisation of the candidate effectors based on expression vector, Apo candidates carry the M. truncatula PR-1 signal peptide for apoplast localisation (Debler et al., 2021). Bottom right, a zoomed image highlights weak necrosis for PnSsp1 and PnSsp17, with white arrows marking necrotic regions.

**Table 3:** Summary of upregulated candidate effectors tested in this study. SnTox3 included for reference. Putative product/structural families were defined by (Jones et al., 2024) or in this study. *Log_2_* Fold Change represents expression change between SN15 and *Δtoxa13* (adj. *p* < 0.0001). “Pangenome candidate” represents whether the candidate effectors were previously predicted (Jones et al., 2024). Predector (Jones et al., 2021) scores of characterised effector and candidates, with a significance cutoff of >2. *EffectorP 3.0* (Sperschneider and Dodds, 2022) predictions of characterised effectors and candidates. Maximal expression of effector genes was determined by (Ipcho et al., 2012). Candidate effector gene presence/absence is based on a *P. nodorum* pangenome by (Jones et al., 2024). “Cyt”/”Apo” – predicted as a cytoplasmic and/or apoplastic effector, respectively. n.d. – not determined.

| Name | Putative product/structural family | $\log_2$ Fold Change | Pangenome candidate | *EffectorP 3.0* predicted | Maximal expression | Gene presence (%) | Wheat symptoms<sup>#</sup> | *N. benthamiana* symptoms<sup>^</sup> |
| --- | --- | --- | --- | --- | --- | --- | --- | --- |
|  |  |  |  |  | *in-planta* |  | *P. pastoris* culture filtrate | *Agrobacterium* transient |
| *SnTox3* | Tox3 NE | - | Yes | Yes (Apo) | Early infection | 92.49 | Necrosis | n.d.* |
| *SnTox267* | Tox267 NE | +3.46 | Yes | No | Early infection | 100 | Necrosis** | n.d. |
| *PnSsp1* | DUF4360-containing | +2.17 | No | Yes (Apo) | Late infection | 100 | No symptoms | Weak necrosis (Cyt) |
| *PnSsp10* | Unknown | +2.64 | No | Yes (Apo) | Early infection | 100 | No symptoms | No activity |
| *PnSsp11* | RALPH family | +1.58 | No | Yes (Cyt/Apo) | Early infection | 100 | No symptoms | Weak necrosis (Cyt) |
| *PnXyn1* | GH11 xylanase | +1.85 | Yes | Yes (Apo) | Early infection | 100 | No symptoms | Necrosis (Apo/Cyt) |
| *PnGlc1* | GH131 glucanase | +1.1 | Yes | Yes (Apo) | Early infection | 100 | No symptoms | Necrosis (Apo) |
| *PnSsp14* | Unknown | +1.07 | No | Yes (Cyt/Apo) | Early infection | 100 | No symptoms | No activity |
| *PnSsp15* | Unknown | +2.12 | No | Yes (Cyt/Apo) | Early infection | 93.64 | No symptoms | No activity |
| *PnSsp16* | Unknown | +1.49 | No | Yes (Apo) | Mid infection | 100 | Inconsistent symptoms | No activity |
| *PnSsp17* | Alt-A1 family | +1.06 | No | Yes (Apo) | Early infection | 100 | No symptoms | Weak necrosis (Cyt) |
| *PnSsp19* | Unknown | +1.13 | No | No | Late infection | 100 | No symptoms | Necrosis (Cyt) |
| *PnSsp20* | DUF3237-containing | +1.42 | No | No | Early infection | 100 | No symptoms | No activity |
| *PnSsp22* | RALPH family | +1.58 | No | Yes (Apo) | Early infection | 98.84 | No symptoms | No activity |
| *PnSsp24* | Unknown | +2.44 | No | Yes (Apo) | n.d. | 100 | No symptoms | No activity |
| *PnSsp25* | SnTox1 homolog | +1.54 | Yes | Yes (Apo) | n.d. | 51.45 | No symptoms | Necrosis (Cyt/Apo) |

#Wheat infiltration images available in **Supplemental Figure S3**.
^Images of transient expression into *N. benthamiana* available in **Figure 4** or **Supplemental Figure S4**.
\*Previous *Agrobacterium*-mediated transient expression does not describe *SnTox3* inducing necrosis in *N. benthamiana* (Dagvadorj and Solomon, 2021).
\*\*SnTox267 induces necrosis when infiltrated into sensitive wheat lines (Richards et al., 2022) (**Supplemental Table S4**).

Structural analysis of the seven necrosis-inducing effector candidates revealed conserved core folds across both bacteria and fungi, as well as within major effector families (Derbyshire and Raffaele, 2023a). *In silico* models of five of these candidates were generated with high confidence (average pLDDT score >90 across the majority of rigid-backbone) superficially matching experimentally-resolved X-ray crystal structures with TM-scores above 0.5 (Zhang and Skolnick, 2005) and alpha-Carbon (Cα) root mean square deviations (RMSD) of less than 5 Å (**Figure 5A**). The effector candidate *PnSsp11* carried a predicted central α-helix surrounded by two β-sheets – conserved amongst *Blumeria graminis* AVR effectors (Cao et al., 2023), while *PnSsp17* carried a putative profilin-like β-barrel fold resembling the *Alternaria alternata* Alt-A1 allergenic effector (Chruszcz et al., 2012) and the *Verticillium dahliae* effector PevD1 (Zhou et al., 2017). The putative glucanase PnGlc1 carries a glycoside hydrolase family 131 catalytic domain structure; β-jelly roll fold flanked by minor α-helices, as observed in the *Coprinopsis cinerea* glucanase CcGH131A (Miyazaki et al., 2013). Structural predictions of PnXyn1 indicate a prototypical xylanase (family 11)-like fold (Hakulinen et al., 2003), albeit with a longer fingers region. The bacterial secreted protein Bd1399 is currently the only experimentally resolved structure of a sole DUF4360 domain protein (Alexander et al., 2023), a domain putatively carried by PnSsp1 and adopting a β-sandwich arrangement that resembles components of diverse CAZyme structures (Cid et al., 2010, Yin et al., 2009). While no experimental structures currently exist for SnTox1, the candidate effector PnSsp25 is predicted to adopt an SnTox1-like fold (**Figure 5B**). The final necrosis-inducing effector PnSsp19 is projected to structurally arrange into an unremarkable single helical motif flanked by long-tailing intrinsically disordered regions, although it should be noted the model quality for PnSsp19 (and PnSsp25) is markedly poorer than that of the other structurally characterised candidate effectors described here.

**Figure 5:**
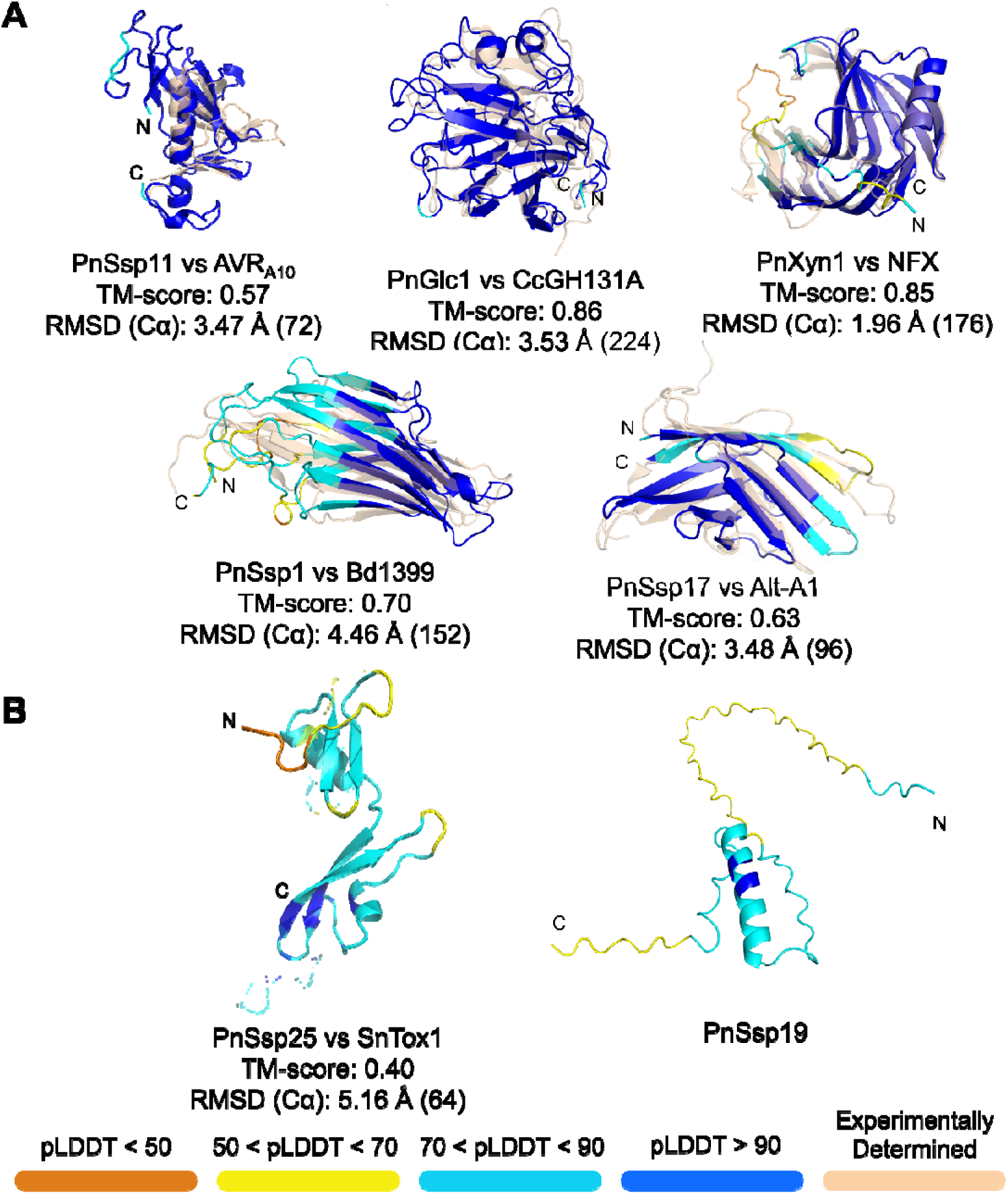
Structural analysis of N. benthamiana necrosis-inducing effector candidates reveals conserved folds. **(A)** AlphaFold 3 *in silico* structural predictions of candidate effectors aligned with experimentally determined representative structures: AVR_A10_ (pdb_00008oxi (Cao et al., 2023)), CcGH131A (pdb_00003w9a (Miyazaki et al., 2013)), NFX (pdb_00001m4w (Hakulinen et al., 2003)), Bd1399 (pdb_00008okh (Alexander et al., 2023)) and Alt-A1 (pdb_00003v0r (Chruszcz et al., 2012)). The experimentally-determined structures (pale orange) in each structural alignment have reduced opacity. **(B)** Predictive structural models of effector candidates PnSsp25 aligned to SnTox1 predicted model (left), and PnSsp19 (right). “N” and “C” show the locations of the N-terminus and C-terminus in the structures, respectively. The predicted models are coloured according to predicted local distance difference test (pLDDT) score per residue; shown at the bottom. TM-align (Zhang and Skolnick, 2005) generated scores (TM-scores) and Root Mean Standard Deviation (RMSD) with the number of alpha carbons (Cα) of the alignment are shown below each model.

## DISCUSSION

Necrotrophic effectors are major determinants of virulence in *P. nodorum* (McDonald et al., 2023, Kariyawasam et al., 2023), yet simultaneous deletion of *SnToxA*, *SnTox1* and *SnTox3* does not significantly reduce pathogenicity on commercial wheat varieties (Phan et al., 2018, Tan et al., 2015). Previous studies showed that removal of individual necrotrophic effector genes can increase the expression of remaining effectors (Phan et al., 2016, Peters Haugrud et al., 2022, Richards et al., 2022). Our work here extends this work and provides a global view of effector expression change in the *Δtoxa13* mutant. This indicated that maintained virulence of *Δtoxa13* is associated with compensatory activation of multiple candidate effectors that exhibit non-host necrosis-inducing activity, transcriptomic epistatic upregulation of the characterised NE *SnTox267*, and subsequent SnTox267-dependent disease phenotype as determined by QTL analysis.

Our host-specific RNA-seq analysis revealed very few transcriptome-specific changes between SN15 wildtype or *Δtoxa13* infection. Furthermore, the host transcriptional response associated with *Δtoxa13* infection is consistent with general responses previously reported following infiltration of the NE SnTox3 and (Ptr)ToxA, particularly the upregulation of primary metabolism, general metabolic processes and redox-associated functions (Winterberg et al., 2014, Pandelova et al., 2009). This overlap supports the view that, despite loss of *SnToxA*, *SnTox1* and *SnTox3*, *Δtoxa13* still induces a host-transcriptome response characteristic of necrotrophic effector activity. The most parsimonious interpretation is therefore that additional necrotrophic effectors compensate for the lack of major NEs, indicating substantial functional redundancy within the *P. nodorum* effector repertoire. Consistent with this, none of the tested candidates exhibited classical NE activity on wheat, except for PnSsp16 which had inconsistent activity, indicating that most function outside canonical host-specific effector interactions. One implication of these findings is that the NE-host interactions all deploy the same strategies to subvert host immune systems for their benefits. Each specific effector-receptor interaction would lead to specific orchestrated defence responses. It is therefore important to identify the triggering molecular mechanisms and understand how they work together to induce disease. This finding also explains the previous observation that the accumulation of numerous necrotrophic effectors in *P. nodorum* may not be explained solely by additive effector-susceptibility-gene interactions (Tan et al., 2015, Phan et al., 2016). Variation in effector content would broaden the range of compatible host genotypes that can be colonised, thereby increasing the persistence of *P. nodorum* as a pathogen on diverse wheat varieties. By contrast, in the wheat tan (syn. yellow leaf) spot pathosystem caused by *Pyrenophora tritici-repentis*, improvement in wheat resistance has been strongly associated with reduced ToxA sensitivity, consistent with a comparatively greater contribution of ToxA to tan spot severity across commercial varieties (See et al., 2024). In addition, recent discoveries surrounding the asymmetric cooperativity of *P. tritici-repentis* and *P. nodorum* during co-infection highlight a potential for diverse effector mechanisms driving fungal pathogenicity progression (Lenzo et al., 2026). Together, these comparisons suggest that *P. nodorum* retains a broader arsenal of partially overlapping virulence determinants, whereas in *P. tritici-repentis* the acquisition of ToxA appears to have had a significant correlation with disease severity of rated commercial wheat varieties.

Unlike *SnToxA*, *SnTox1* or *SnTox3*, which exhibit a high of presence/absence variation in worldwide *P. nodorum* populations (McDonald et al., 2013), *SnTox267* is absent from only a small subset (<5%) of *P. nodorum* genomes (Richards et al., 2022) and is carried in 100% of 173 previously defined *P. nodorum* isolates (Jones et al., 2024). SnTox267 as a NE is characterised to infect susceptible wheat lines carrying the S-genes *Snn2* and *Snn6* cooperatively, or *Snn7* in a light-independent manner (Richards et al., 2022). Our results here demonstrate SnTox267-specific S-genes are carried on just over half of tested Australian elite commercial cultivars, reflecting the need to accommodate programs to combat SnTox267-sensitivity within *P. nodorum* isolates. Furthermore, the high frequency of SnTox267 sensitivity in commercial lines is a likely contributing factor for the maintenance of *Δtoxa13* virulence that was previously observed (Phan et al., 2018).

Non-host effector activity has been well-characterised in the wheat pathogen *Zymoseptoria tritici,* causal agent of septoria tritici blotch. *Z. tritici* retains a collection of diverse, secreted effectors to suppress host immunity and facilitate host infection, yet these effectors are also able to induce cell death in the non-host *N. benthamiana* (Welch et al., 2022, Thynne et al., 2024, Kettles et al., 2017). Six of the seven identified candidate effectors identified in this study which induced necrosis did so when expressed without a secretion signal, and four of these were exclusively cytoplasmic, suggesting dependence on wheat-or monocot-specific external receptors or internalisation processes absent in the *N. benthamiana* heterologous system. The apparent inactivity of these candidate effectors during wheat infiltration may instead reflect a requirement for fungal-mediated delivery or host-specific uptake mechanisms that are not recapitulated in isolation. However, we cannot exclude the possibility that the necrosis observed in *N. benthamiana* reflects activation of conserved dicot immune responses, including pattern-triggered immunity elicited by microbial-associated molecular patterns or cell wall-degrading enzyme activities, rather than a direct contribution of these candidates to virulence in wheat. Additionally, these candidate effectors may instead have some suppressor-mediated epistatic function, analogous to those observed in other fungal systems such as powdery mildew, where effector-induced cell death is actively repressed despite the presence of a cognate avirulence factor (Bernasconi et al., 2026).

Structural classification further highlights the diversity of the candidate non-host effector repertoire. Five *P. nodorum* effector candidates identified in this study fall within major fungal effector families defined across 20 species (Derbyshire and Raffaele, 2023a). PnSsp11 and PnSsp22 are predicted to adopt RALPH-like folds (**Figure 5A**), a family best known from cereal powdery mildew avirulence proteins (Cao et al., 2023, Praz et al., 2017). The presence of RALPH-like analogues in a necrotroph suggests that related effector scaffolds can be repurposed for contrasting pathogenic strategies, as illustrated by the structural similarity but functional divergence of Avr2 and ToxA (Di et al., 2017, Faris et al., 2010, Sarma et al., 2005). PnGlc1 is an apoplastic effector belonging to glycoside hydrolase family 131 (GH131). GH131 glucanases have broad substrate specificity and are characteristic of plant-colonising fungi and oomycetes (Anasontzis et al., 2019). However, their capacity to induce necrosis has not been previously demonstrated. In phytopathogenic fungi, GH131 proteins such as PnGlc1 may therefore represent an effector-adapted variant of this enzyme class. The Alt-A1-like predicted β-barrel fold in PnSsp17 is also observed in characterised effectors, including PevD1 from the broad-host fungal pathogen *Verticillium dahliae* which induces necrosis in *N. benthamiana* (Zhou et al., 2017) and is upregulated during host infection (Zhang et al., 2019). The same β-barrel fold is also present in necrosis-and ethylene-inducing peptide 1 (Nep1)-like proteins, which induce necrotic symptoms in *N. tabacum* and *Arabidopsis* hosts (Ottmann et al., 2009).

The *SnTox1*-like candidate effector PnSsp25 may have emerged from gene duplication and may predate speciation from *P. pseudonodorum*; which carries both *SnTox1*-and *PnSsp25*-like genes (**Supplemental Figure S5**). Given that recognition residues can vary substantially within a shared structural family (Lazar et al., 2022, Cao et al., 2023), duplication may have facilitated diversification of host targets in *P. nodorum*, something observed for SnTox3 and the SN15-absent NE SnTox5 (Kariyawasam et al., 2022). Seven GH11 xylanases are encoded within *P. nodorum* SN15, although each appears to have diverged before speciation, as each paralogue groups more closely with orthologues from related species than with other *P. nodorum* copies (**Supplemental Figure S6A**). Within the GH11 family, PnXyn1 is notable in behaving as cytoplasmic non-host effector that induce necrosis in *N. benthamiana* without apoplastic targeting. CmXyn1, which shares high amino acid identity with PnXyn1 (**Supplemental Figure S6B**), has been described as an apoplastic effector, although assays without a signal peptide gave inconsistent results (Lee et al., 2023). PnXyn1 and CmXyn1 may therefore define a distinct functional subgroup within fungal GH11 xylanases. PnSsp1, a necrosis-inducing non-host effector, is predicted to contain DUF4360. This domain also occurs in the *Verticillium dahliae* non-host effector VdSCP27, which induces necrosis in *N. benthamiana* (Wang et al., 2020). Genes encoding secreted DUF4360 proteins are strongly induced during plant-associated growth in multiple fungi, including *Colletotrichum lindemuthianum* and the symbiont *Hebeloma cylindrosporum* (de Queiroz et al., 2019, Doré et al., 2017), and may therefore represent a secretion-adapted protein class broadly used by plant-colonising fungi to modulate plant interactions. Future work, leveraging structural bioinformatics (Verdonk et al., 2025, Jones and Raffaele, 2025), may enable direct identification of the biological targets for these structurally conserved effector candidates from *P. nodorum*.

In fungi, long promoters are commonly associated with stress-responsive genes and more complex regulatory inputs (Kristiansson et al., 2009). The enrichment of long promoters among genes upregulated in *Δtoxa13 in-planta*, including candidate effectors, is therefore consistent with an adaptive transcriptional response to the loss of major necrotrophic effectors. This premise supports a relationship between intergenic length and regulatory plasticity rather than neutral genome drift (Noble and Andrianopoulos, 2013). Effector regulation by *P. nodorum* transcription factors, including the archetypal *SnToxA* and *SnTox3* regulator PnPf2 (Rybak et al., 2017, John et al., 2024), is hypothesised to be linked to chromatin accessibility (Morikawa et al., 2026). As host-induced chromatin remodelling is highly dynamic in other fungal pathogens (Gay et al., 2021, Kramer et al., 2023), direct tests such as histone occupancy profiling or targeted disruption of chromatin modifiers (Clairet et al., 2024, Connolly et al., 2013, Meng et al., 2021, Zhang et al., 2021) should help clarify whether similar mechanisms operate in *P. nodorum* for regulation of candidate effector genes.

While non-host effectors identified here are not directly linked to wheat-specific virulence, their conservation and activity across divergent plant systems suggest they may represent a broader class of latent virulence factors shared among cereal pathogens. This study may posit two broad inferences regarding the pathogenic biology of *P. nodorum*: First, functional redundancy among necrotrophic effectors may help explain how this highly specialised wheat pathogen persists across host populations with diverse genotypes-gene profiles. Population-level surveys have shown substantial variation in effector gene content among *P. nodorum* isolates, including strains lacking one or more of the major characterised effectors yet remaining pathogenic on wheat (McDonald, 2025, McDonald et al., 2013, Stukenbrock et al., 2006). Such variation is consistent with a model in which alternative effectors can substitute for one another to maintain virulence, thereby allowing infection of wheat lines that differ in their complement of S-genes.

Second, the detection of multiple non-host effectors suggests an additional layer of host specialisation in *P. nodorum*. If some virulence determinants require wheat-specific uptake, receptors or intracellular compatibility factors, then their activity would be expected to remain restricted to a narrow host range. This interpretation is consistent with broader comparative analyses in which *P. nodorum* is regarded as a narrow-host-range polymertroph on the basis of CAZyme content (Hane et al., 2020), and it may help explain why the species has evolved as a highly specialised cereal pathogen rather than as a broad-host-range necrotroph. This hypothesis is further supported by the QTL detected in *Δtoxa13* which could not be assigned to known loci or sensitivity genes (Phan et al., 2016, Phan et al., 2018), raising the possibility that further specialised host-pathogen interactions are normally masked by the NEs SnToxA, SnTox1 and SnTox3. This further solidifies *P. nodorum* adaption as a specialised pathogen, with dominant necrotrophic effectors suppressing non-host effector activity, while the conservation of these elements implies a retained, context-dependent role in pathogenicity. Collectively, our observations suggest that targeting conserved effector regulatory networks, rather than individual host-specific interactions, may offer a more tractable route to resistance in pathosystems where NE redundancy and compensatory regulation undermine the durability of S-gene removal for plant disease resistance approaches.

## DISCLOSURE OF AI USE

The authors used AI to improve the clarity and readability of the manuscript and subsequently reviewed and edited all content.

## DATA AVAILABILITY

RNA Sequencing data can be found under NCBI Bioprojects PRJNA632579 and PRJNA1479129.

## SUPPLEMENTARY DATA

A supplementary data file is provided with this submission: SUPPLEMENTAL_DATA_Toxa13_JXB.docx

Another file, Supplemental_File_S1.xlsx (45 MB), is available with the RNA-seq differential-expression data from the authors. The JXB online submission portal was unable to convert this document to a PDF format.

## SUPPLEMENTARY DATA FIGURE CAPTIONS

**Supplemental Table S1**: Fungal family, species and strains used for the phylogenetic analyses of GH11 xylanases.

**Supplemental Table S2:** Primers and synthetic DNA used in this study.

**Supplemental Table S3:** Candidate effectors up-regulated in *Δtoxa13* relative to SN15. All candidates are predicted to be secreted using SignalP 6.0 (Teufel et al. 2022) and being >80 amino acids (AA) and <350 AA in size. Average normalised transcripts for SN15 and *Δtoxa13* are for *in-planta* RNA-seq dataset.

**Supplemental Figure S1:** Principal component analysis of transcriptomes of wheat-host against SN15 and *Δtoxa13* infection three-days post-inoculation, and uninfected leaves (tween-treated). Three biological replicates were used for Tween-treated leaves and *Δtoxa13*-infected (this study), and four biological replicates were used for SN15-infected leaves (Jones et al. 2019).

**Supplemental Figure S2:** Correlations in the upstream intergenic region (UIR) of all differentially regulated genes in *Δtoxa13 in-planta*. Distribution density plot of UIR lengths between down-regulated (Down), up-regulated (Up) and non-differentially regulated (Same) genes in *Δtoxa13 in-planta*, with the median lengths marked with a vertical dotted line. The median lengths significantly differ between groups according to the Kruskal-Wallis test and the *post-hoc* Dunn’s test with the Holm adjustment (adj. *p <* 0.001).

**Supplemental Figure S3:** *Pichia pastoris* culture filtrate infiltration of up-regulated candidate effectors from *P. nodorum Δtoxa13* into commercial wheat cv. Halberd, Axe, Emu Rock, Mace; seven days post infiltration.

**Supplemental Figure S4:** Agroinfiltration of negative/non-host necrosis inducing up-regulated candidate effectors from *P. nodorum Δtoxa13* and expressed in SN15. Necrosis inducing effector candidates are shown in-text at **Figure 4**. “White light” shows representative *Nicotiana benthamiana* leaves agroinfiltrated with one of 14 candidate effectors and photographed 10 days post-infiltration. NLP2 (necrosis-and ethylene-inducing peptide 1 (NEP1)-like protein 2) and GFP (green fluorescent protein) are the positive and negative controls, respectively. Black names show no necrosis-inducing activity. The location of each agroinfiltration has been partially randomised. “Necrosis/GFP” shows the same leaf highlighting necrosis (pink) and GFP fluorescence (green). “Cyt” and “Apo” indicate cytoplasmic and apoplastic localisation of the candidate effectors, respectively.

**Supplemental Table S4:** Recombinant expression of SnTox267 infiltration into 78 different varieties of Australian commercial wheat cultivars. A score of 0 indicates insensitivity (white, no reaction); 1, slight chlorosis (green); 2, moderate chlorosis/slight necrosis (yellow); 3, moderate necrosis (orange); 4, extensive necrosis (red). (*one replicate of Baxter unsuccessful).

**Supplemental Figure S5:** Sequence analysis of candidate effectors PnSsp25. A protein sequence alignment of PnSsp25/SnTox1 homologs in *P. nodorum* and *P*. *pseudonodorum.* “SignalP” is the predicted signal peptide. The purple bar represents the location of the putative Kex2 cleavage site based on SnTox1/PnSsp25, and orange arrows indicate the location of 16 conserved cysteine residues in SnTox1 and PnSsp25. A phylogenetic tree showing the relatedness of PnSsp25/SnTox1 homologs in *P. nodorum* and *P*. *pseudonodorum*. The percentages next to the clades show the percentage sequence identity within the clades.

**Supplemental Figure S6:** Phylogenetic analyses and characterisation of the GH11 xylanase PnXyn1. (**A**) A phylogenetic tree showing GH11 xylanases in 23 fungal species, three non-fungal species and a necrosis-inducing GH10 xylanase (ppxyn1) as an outgroup. The nodes are coloured according to fungal classes. Red circles next to the nodes indicate xylanases required for full virulence by the fungi of origin. Orange circles indicate a necrosis-inducing activity on the host and/or non-host plant. PnXyn1 and the closely related PtrXyn1 are bolded. Grey circles indicate that the fungi of origin are non-pathogens. The scale bar indicates amino acid changes per site. (**B**) Sequence alignment of 22 GH11 xylanases, including structurally determined GH11 xylanases XynC (PDB: 1BK1) and XYN1 (PDB: 1XYN), showing putative N-terminal Kex2 cleavage sites only conserved in fungi. Grey represents predicted signal peptides. Putative Kex2 cleavage sites are coloured according to their recognition site – blue ((K/R)R), pink (LXXR) and purple (LX(K/R)R). The size differences of the putative Kex2 cleavage sites are due to gaps in the sequence alignment within the sites. The dotted box represents conservation of putative Kex2 cleavage sites in the N-termini of fungal GH11 xylanases. Red represents regions required or sufficient for necrosis-inducing activity on tested *Nicotiana* spp. The orange bars represent structurally determined regions of XynC and XYN1. The numbers show the length of the protein alignment. The brown and black bars on the right encompass GH11 xylanases of fungal and bacterial origin, respectively. The black triangles at the top of the alignment show the location of the conserved catalytic glutamic acid residues.

## AUTHOR CONTRIBUTIONS

Shota Morikawa: Investigation, Conceptualisation, Formal analysis, Data curation, Methodology, Validation, Writing - original draft. Callum Verdonk: Investigation, Validation, Supervision, Formal analysis, Writing - original draft, Writing - review & editing. Leon Lenzo: Data curation, Methodology, Formal analysis. Huyen Phan: Investigation, Methodology, Formal analysis. Eiko Furuki: Investigation, Methodology. Kristina Gagalova: Formal analysis. Johannes Debler: Methodology. Chala Turo: Resources. Carl Mousley: Supervision. Evan John: Supervision. Bernadette Henares: Supervision, Methodology. Kar-Chun Tan: Conceptualisation, Project administration, Resources, Supervision, Funding acquisition, Writing - review & editing. All authors contributed to finalised manuscript prior to submission.

## ACKNOWLEDGEMENTS

The authors thank Darcy Jones, Kasia Clarke, Christina Grimes and Julie Lawrence for their assistance during the experimental undertakings. We thank the Analytics for the Australian Grain Industry (AAGI) for the advice on the RNAseq data processing and data standards.

This study was conducted by the Centre for Crop and Disease Management, a co-investment between the Grains Research and Development Corporation (GRDC) and Curtin University – grant CUR1403-002BLX. The data management was supported by AAGI – grant CUR2210-005OPX. SM was supported by an Australian Government Research Training Program Scholarship administered through Curtin University. The funders had no role in study design, data collection and analysis, decision to publish, or preparation of the manuscript.

