## Supplemental Data for "Major effector loss reveals compensatory pathogenicity networks in a necrotrophic wheat pathogen"

†These authors contributed equally.

**Supplemental File S1** (separate .xlsx file) contains RNA-Seq data analysis (differential-expression and gene-ontology term enrichment) for *in planta* and *in vitro* SN15 and *Δtoxa13* fungal reads, as well as wheat-host transcriptome to SN15 and *Δtoxa13* infection and Tween-treated samples. RNA reads can be found under NCBI Bioproject PRJNA632579 (SN15) and PRJNA1479129 (*Δtoxa13* and Tween).

**Contents**

**Supplemental Table S1	2**

**Supplemental Table S2	3**

**Supplemental Table S3	6**

**Supplemental Figure S1	9**

**Supplemental Figure S2	10**

**Supplemental Figure S3	11**

**Supplemental Figure S4	12**

**Supplemental Table S4	13**

**Supplemental Figure S5	14**

**Supplemental Figure S6	15**

**Supplemental Table S1**: Fungal family, species and strains used for the phylogenetic analyses of GH11 xylanases.

| Fungal class | Species | Strains | Classification |
| --- | --- | --- | --- |
| Dothideomycetes | *Alternaria alternata* | SRC1IrK2f | Pathogen |
|  | *Ascochyta lentis* | Al4 | Pathogen |
|  | *Ascochyta rabiei* | ME14 | Pathogen |
|  | *Bipolaris sorokiniana* | ND90r | Pathogen |
|  | *Colletotrichum graminicola* | M1.001 | Pathogen |
|  | *Leptosphaeria maculans* | CAN1 | Pathogen |
|  | *Parastagonospora nodorum* | SN15 | Pathogen |
|  | *Pyrenophora tritici-repentis* | M4 | Pathogen |
|  | *Pyrenophora teres* f. *maculata* | SG1 | Pathogen |
|  | *Pyrenophora teres* f. *teres* | W1-1 | Pathogen |
|  | *Zymoseptoria tritici* | IPO323 | Pathogen |
| Sodariomycetes | *Fusarium graminearum* | PH-1 | Pathogen |
|  | *Fusarium oxysporum* | Fo47 | Non-pathogen |
|  | *Magnaporthe oryzae* | 70-15 | Pathogen |
|  | *Neurospora crassa* | OR74A | Non-pathogen |
|  | *Trichoderma reesei* | QM6a | Non-pathogen |
|  | *Verticillium dahliae* | Vd991, VDG1, VdLs.17 | Pathogen |
| Leotiomycetes | *Botrytis cinerea* | B05.10 | Pathogen |
|  | *Sclerotinia sclerotiorum* | 1980 | Pathogen |
|  | *Valsa mali* | LXS080901 | Pathogen |
| Eurotiomycetes | *Aspergillus flavus* | NRRL3357 | Pathogen |
|  | *Aspergillus nidulans* | FGSC A4 | Non-pathogen |
| Ustilaginomycetes | *Ustilago maydis* | UM521 | Pathogen |

**Supplemental Table S2:** Primers and synthetic DNA used in this study.

| **Primer Name** | **Sequence** |
| --- | --- |
| Tox267_3R | AGCAAACATGGCAGATGTCG |
| Tox267_3R2 | AGATAGGATCGAAAGCAGCC |
| Tox267_3Screen_R | AATTGTTGATCTGATCGGGG |
| Tox267_5F | GCATTGAGACTAGGTACAGG |
| Tox267_5F2 | CTTTTTCCTGCTCTTCCTCG |
| Tox267_5Screen_F | GAAGATCGGATGCCACCTAG |
| Tox267KO_pgpdA_5R | AGAGCTCACGAGTTCGTCACATTTTTCTACGCTTGCAGCAAGGTGTAGTG |
| Tox267KO_tTrpC_3F | TACAACTCTCCTATGAGTCGTTTACCCAGAGATCAGGATCTGCATCAGG |
| M13_Gateway_F | GTAAAACGACGGCCAGT |
| M13_Gateway_R | CAGGAAACAGCTATGAC |
| JD029_pDEST1_F | AAAACCGCTCACCAAACATA |
| JD030_pDEST1_R | TTTTCTTTGAAACAGAGTTTTCC |

| **Synthetic DNA Name** | **Sequence** |
| --- | --- |
| SNOG_00075 GATEWAY  (PnSsp1) | GGGGACAAGTTTGTACAAAAAAGCAGGCTCCATGCTCCCAGGCCTCCCCACCGTCGAGCTCGGTGAAGCTCCTCCGGCCGGCTCGGTCACCATCAAGGGCGTCAGCTATGGAGGAACTGGATGTCCCCAGGGCACCATGAGCTCCCAGATCTCTTCGGACCGCACCATTGTCACCCTCATCTTTGACTCTTACATCGCTTCCACCGGCCCTGGCATCTCTGTCACGGAGCAGCGCAAGAACTGCCAGCTCAACGTTGACCTTCAGTACCCCGGAGGCTTCCAGTACTCCATCTTGTCCGCCGACTACCGCGGTTACGCCGCCATCCAGAAGGGCATCACGGGCACTCTCAAGTCGACCTACTACTTCTCTGGCCAGACTGCTCAGACGTCCACCGAGTACAACTTCGTAGGCCCCGTCAACGGCGACTACCTCAAGCACGACGAGGCCGACTCGACCTCCATCATCTGGTCGCCCTGCGGCGCCGCCGGCATGCTCAACATCAACTCGCAGGTGCGCCTGACCAGCACCAACTCGTCGGCGACCGGTCTGCTCACCACCGACTCGACCGATCTCAAGTTCAGCCAAGTCGTGTACGTCCAGTGGCAGAAGTGCACCAAATACCCAGCTTTCTTGTACAAAGTGGTCCCC |
| SNOG_07039 GATEWAY  (PnSsp10) | GGGGACAAGTTTGTACAAAAAAGCAGGCTCCATGCCAGCCGAGCTCCAGGAGCGCCAATGTATTTTCAACGGATTGGGTTGCGACGTTTTGAATAATCGATGCTGCTCACGTAATTGCGAATCTTCTGTATGCGGAGGTCGGACCTTTACATGCCAGCCTCAAACTACCGGATGCTCCCCGCCAGTGCCAGGGTCATACCCAGCTTTCTTGTACAAAGTGGTCCCC |
| SNOG_08125 GATEWAY  (PnSsp11) | GGGGACAAGTTTGTACAAAAAAGCAGGCTCCATGCTCTGGAACCAGGAGCTTTGTAAAGGCGCCGGTGCATGCATAGACATAGGTGTCTTGGTCTCATATCCCTTTCGCTGTCCCGATGGTAGCGCAATCACGAGACCTTCCTTCGGGACGGATTTGCAAAAAGCTGCTCTGGCCGGAGCTACTCGAATTACCAAAGAAGAGTTTCCAAAGACGTGTTACGCGGGCAAAGTTCCCAGCGCGAATGCCATATTTGTTCGCACCACGACGCGCAACGGTCAAACTGCCTATACCTTCATCGAAGAGGGTTGTACCGACCCGAACCCAAAGTTCCCTAAGGATTGCTACCATTACACAACCAACGCATCTACATACACATTCTGCCAGCTAGTCGATGCGAAAGGTGGACAGTGTACCGAGAACCTACAAGCCGGTAGGTGTGAGCGTTGGGGTGATACAGCTGGCCGGACCGAGTGCAAAGATTGGAAGATTGGCCAGCCTGATTTTCCCGATGATGAGTACCCAGCTTTCTTGTACAAAGTGGTCCCC |
| SNOG_09650 GATEWAY  (PnXyn1) | GGGGACAAGTTTGTACAAAAAAGCAGGCTCCATGGATGCTCCTGATTTCGAGCTGACCGCCAGCAACATTGTTCGTCGCCAGGACTACAACCAGAACTACAAGACCAGCGGCAACGTCAACTTCCAGCCCACCAACAACGGCTACTCTGTCCAGTTCTCCAACGCTGGTGACTTCGTTGTCGGAAAGGGATGGAAGACCGGAAGGGACCGCAAGATCAACTTCAGCGGTTCTACTTCCGCCACTGCCGGCACGGTCCTCGTCTCCATCTACGGATGGACCACCTACCCTCTCGTTGAGTACTACATTCAGGAGTACACCAGCAACGGCGCTGGCTCTGCTCAGGGTCAGAAGATGGGCACGGTCACCTGCGACGGCTCCGTCTACGACATCTGGAAGCACACCCAGACCAACCAGCCTTCCATCCAGGGTACCTCCACCTTCCCCCAGTACATCAGCAACCGTCGGACAAAGCGCCCCGGCAGCGGCACCGTCACCACCAAGTGCCACTTCGACGCCTGGGCCAAGCTCGGCATGAAGCTCGGTACCCACAACTACCAGACCCTTTCCACTGAGGGCTGGGGCAACGCCGGCGGTAACTCCAAGTACACCGTCTCCGGCTCTTACCCAGCTTTCTTGTACAAAGTGGTCCCC |
| SNOG_10382 GATEWAY  (PnGlc1) | GGGGACAAGTTTGTACAAAAAAGCAGGCTCCATGGCCGAAGTCAAGTGCACCGTCGTCTTCGATGGTCGTGTCCCGGTCAATAGCACACCCACCTCTTTCGATACCAGCAACGCTTTGTTCAACCCCGACTACGTCAAGGGCAACAACTTGACTTGGAGCCAGATCCTCCAGTTCCCCAAGGACGCCTCTCGCTTTGACGGTAACAAGTTCAAGGCTGTCGAGGTTACCATCTCGGACAAGTCGATCTTCCAGAAGCAGAATGGATTCCGCCGCGCAGGCCTCCAGTTCGCCAAGGACGCGCCTGATGGCGAGGGCGGCAAGGGTGTCAAGACGCTACACTGGAGCGTAAAGCAGGACCAGGCTAGGCCCTTGAACCTGACCCACGAGTACCTCAACGTGTGGCACGAGACTGCCGACTACTCAGCGAACCAGATCCAGTTCCAGACAGGCTCGCTCATCGGCAAGTCTGACGCAGACAAGAGCAACTTCAAGATTCTTGACCGCAGCGGCAACTTCCTGTGGTCTGTTGGTATCGACCAGAGGAACTGGCAGAACTTCGCCGTCAAGCTTGACTACAACCAGAACCAAGTTTCCATCTACTACTCCATTGGAAACTCCGAGCTGTCCGAAGTCGTGACCAACAAGACCGTCAACATTGCCGGAGGTGGCCAATTCCAGCTCGGCATGCTCAAGAAGCCCACTGGAACTTCCGATGTGGCCAACGCTGGTTACCAGTCTCCTAACCTCAACGAGGGCCAGATTTACGGTGGTATCTTCCTCGAGGACTCTGCCAACGGCTGTGTCTCCCGCTACCCAGCTTTCTTGTACAAAGTGGTCCCC |
| SNOG_11453 GATEWAY  (PnSsp15) | GGGGACAAGTTTGTACAAAAAAGCAGGCTCCATGCCTGCAGAGCTTCAGGAGCGACAATGTCAAGGCCGTGGCGGTCGCTGCGCCGGGCAATACGAATGCTGCGCCAATACCTACTGTGTTGAAATTATTTGTAACTCGAGCAAATTTTTTTGCGAGCCTCTTGGCGCCACTCGTTGTATAAACCCCGGCAGCGGCGGCGGCCGCCATTACCCAGCTTTCTTGTACAAAGTGGTCCCC |
| SNOG_12448 GATEWAY  (PnSsp16) | GGGGACAAGTTTGTACAAAAAAGCAGGCTCCATGACACCAACTCCACAGCGACCTGTCCCTGGCGCACCAGCTGACCCTGGCGTCAGAGTGCCCAAGGGCACTTTCTGTACCTTCACCTCGCAAGAAGGCGTCCTAGGATGCAATACTGGCGATGGCGGCACATTCAACATCATCGACGGAAAGATTAGCGGCTGTGCTGGATGCACGAAGGAGAACGGATTCGGAAAATACCCAGCTTTCTTGTACAAAGTGGTCCCC |
| SNOG_12449 GATEWAY  (PnSsp17) | GGGGACAAGTTTGTACAAAAAAGCAGGCTCCATGGAGAACGTCACCATCTCCAACTTCCTCTACGTCGGTGTCAGCGGCTACGACCAGATCTCTTTCAGCCTATCCGTGGACGACATCAACTGCGGCGCTGACCACTACGTGATTGGTGGTATGTACGCCTGCGACAACAAGGCGTGGACCTTCCAGATCAACGAGGCACAAGGCCACCAAATCAAGTTGCTACACGCTGTCAACGGCAAAACTCTTTCCGGCGACTTCGACATCAAGATGAACGGTCCTATCACAACTGTGAGACAGCAGATTGGTACTTCCACAGCCGAATTGAACTACCCAGCTTTCTTGTACAAAGTGGTCCCC |
| SNOG_13405 GATEWAY  (PnSsp19) | GGGGACAAGTTTGTACAAAAAAGCAGGCTCCATGAAGCCGTGGCCGTGGGCTGCGCCGGCGGCTAGCCTTGTACCGACTGCAAATGCGCAGCCGAACAAGACACATCGAGGCAACAGGACGAAACATCATCACCACGAGCACACGCCAACGTTCAAAGAGCCGTGCAACTGCCCGCAGCCGATCGTACCTATGAATCTACTTAGCGAGAACGAGAAATGTTTAATGAAGCATGCAGCAGCCATTGGATGTTATATCGGATCGAAGGGCGGATGTCCATCACCGGCACCGGCTTGCGGACTTGGCGTTACCAAAGCGATTCCCTACACTCATTACCCAGCTTTCTTGTACAAAGTGGTCCCC |
| SNOG_14242 GATEWAY  (PnSsp20) | GGGGACAAGTTTGTACAAAAAAGCAGGCTCCATGTACAATGCGCCGAAGCCGCCGACGTTGACCTTATTGTACAGCATGGCATGCGACCTGGCTCCGGAAATGTCAATGGGCGCCGTGCCTACTGGCCAAGAACGCATTATCATTCCCATAATTGGCGGTACATTCAACGGATCCCGCATATCCGGAAAGATTCTGAACGTCGGTGCCGACTGGTACCTCGTGGACAGCCGGGGGAAAGGTCGTCCTGATACACGGTACAATCTCCAAACAAATGACGGAACCTACATATACGTGCAGACAGAGGGGCCGACTTTTGAAGATGGACGGACTCTGCTTCGTGCCAAGTTCGAAGCGCCCATCAATTCGACATACTCGTGGATGAATGAGGTTGTTGGACTGGGGGTGCTGACCGTCAATGGTACGCAGCAAGTGCTGATCGATATGTGGCATGCATCTCTGTACCCAGCTTTCTTGTACAAAGTGGTCCCC |
| SNOG_16438 GATEWAY  (PnSsp22) | GGGGACAAGTTTGTACAAAAAAGCAGGCTCCATGCAAGCCGGCGATCTGTACCGCCAGTTCCCCGAAAGCCTCGACTGCCCGGTGAAGAACGGCGTCCACGTATTGAAGAAGGACCTCGTAGAGGCAGTCAAGAACGGCAAGAGGGACGGACCGCCATATGAGGCAAGTGCGGCAAACCTCGCGACTAGGCATTGTGGAAACTCGAACTTCAAGGGAATCCCTCTCTGGACCACCGAGATTCCAGATGGTACAGAATCCGCTGGTGCTCTTTTCTACGCTGCGGCATCCAACGGCACTTTCTATTTCTGCGGCACAACCTCGGGCCGAGTACCGTCGGGCTGGCCTTCTTCATGCACGGAGAACTACTACCCAGCTTTCTTGTACAAAGTGGTCCCC |
| SNOG_30359 GATEWAY  (PnSsp24) | GGGGACAAGTTTGTACAAAAAAGCAGGCTCCATGGCCCCTCTCGATACCCCTGCCAATGCCGAACTAGTAGCAAGACAGAAATACCCAATTCATCAGGTATCCGAGATATGCTGCTCTAAAGATCATCTAGCGGAGAGGTTCTTCTGCGCTTGGTCGTATCCTGCCACTGGATGCAATCCGTGCCCCTCAGATATCCCTATATGCACAAAGCCTTACCCAGCTTTCTTGTACAAAGTGGTCCCC |
| SNOG_42342 GATEWAY  (PnSsp25) | GGGGACAAGTTTGTACAAAAAAGCAGGCTCCATGAACAAAGAAATCCATACATTCGACAGCCTTGGTCTTACCCGCCGACAGGGTCAACAAGGTGGAATCTGCCATAGTATTGAGGGCGGAGGCTGCCGAGCATACAGTACCAAGGGTCATTGCTGCTGCACAGATTGTAGCAACGTGGAATGCAACGAGGTCTGCACCAACATTAAGCCGACAGAAATTTGTGCGACGTGCTGCAACCGTGGTGGGGAAGCTCACGACGTCTGCTGCCCAGTCGGACTACATGAGAGTCCCTGCGACCCTTGCAAGTCGGCTGGCGCCGGACTGGCACACTGCTACCCAGCTTTCTTGTACAAAGTGGTCCCC |

**Supplemental Table S3:** Candidate effectors up-regulated in *Δtoxa13* relative to SN15. All candidates are predicted to be secreted using SignalP 6.0 (Teufel et al. 2022) and being >80 amino acids (AA) and <350 AA in size. Average normalised transcripts for SN15 and *Δtoxa13* are for *in-planta* RNA-seq dataset.

| **In-text name** | **Locus ID/Gene** | **AA Sequence Length** | **Deeploc Extracellular** | **Log-Fold_2_ Change** | **Adjusted *P-*value** | **Average normalised transcripts SN15** | **Average normalised transcripts *Δtoxa13*** | **Putative product** |
| --- | --- | --- | --- | --- | --- | --- | --- | --- |
| PnSsp11 | SNOG_08125 | 176 | 0.986 | 1.581757 | 7.90E-26 | 11244.28 | 34178.84 | RALPH-like |
| PnSsp22 | SNOG_16438 | 138 | 0.9933 | 1.580937 | 6.23E-28 | 10987.39 | 33366.82 | RALPH-like |
| PnSsp16 | SNOG_12448 | 83 | 0.9999 | 1.486973 | 5.76E-30 | 9001.446 | 25689.91 | Unknown |
| PnGlc1 | SNOG_10382 | 284 | 0.9222 | 1.102936 | 5.32E-11 | 6542.838 | 14405.33 | GH131 hydrolase |
| PnSsp15 | SNOG_11453 | 80 | 1 | 2.121437 | 4.75E-15 | 3050.851 | 13797 | Unknown |
| PnXyn1 | SNOG_09650 | 221 | 0.9997 | 1.850982 | 3.77E-08 | 2519.856 | 9781.098 | GH11 xylanase |
| PnSsp10 | SNOG_07039 | 98 | 0.9946 | 2.644482 | 3.04E-53 | 1469.3 | 9301.082 | Unknown |
|  | SNOG_04279 | 102 | 0.9865 | 2.761305 | 1.52E-57 | 1343.281 | 9212.997 | Unknown |
| PnSsp24 | SNOG_30359 | 86 | 1 | 2.445555 | 5.55E-35 | 1460.556 | 8089.661 | Unknown |
| SnTox267 | SNOG_14493 | 265 | 0.9809 | 3.463461 | 3.08E-41 | 644.5513 | 7277.009 | SnTox267 NE |
| PnSsp17 | SNOG_12449 | 113 | 0.9496 | 1.063078 | 4.19E-13 | 3359.465 | 7152.562 | Cyanovirin-N domain/Alt-A1-like |
| PnSsp20 | SNOG_14242 | 161 | 0.8309 | 1.418454 | 1.67E-08 | 2059.952 | 5773.342 | UPF0311/DUF3237 |
| PnSsp19 | SNOG_13405 | 115 | 0.9995 | 1.133867 | 4.96E-18 | 2013.644 | 4480.759 | Unknown |
| PnSsp1 | SNOG_00075 | 208 | 0.9818 | 2.175796 | 4.61E-28 | 918.1014 | 4225.422 | DUF4360 |
| PnSsp25 | SNOG_42342 | 124 | 1 | 1.543685 | 2.02E-14 | 1403.215 | 4201.083 | SnTox1 paralog |
|  | SNOG_13486 | 161 | 0.9954 | 1.347551 | 1.43E-20 | 1547.67 | 4018.769 | Unknown |
|  | SNOG_11452 | 80 | 1 | 1.073739 | 2.72E-10 | 1714.549 | 3698.918 | Granulins domain |
|  | SNOG_12911 | 118 | 0.9998 | 1.644872 | 3.20E-34 | 1131.205 | 3587.548 | Unknown |
|  | SNOG_04743 | 275 | 0.799 | 1.105192 | 1.62E-18 | 1491.824 | 3262.668 | Alkaline sensitive linkage |
|  | SNOG_44384 | 157 | 0.5926 | 7.666589 | 1.35E-21 | 9.171256 | 2067.15 | Unknown |
|  | SNOG_15451 | 303 | 0.8802 | 1.076354 | 7.70E-08 | 783.8221 | 1705.11 | Carboxylic ester hydrolase |
|  | SNOG_02930 | 244 | 0.9991 | 1.070428 | 4.55E-09 | 778.6139 | 1674.462 | lytic cellulose monooxygenase |
|  | SNOG_06015 | 199 | 0.872 | 1.895864 | 1.72E-14 | 404.5844 | 1553.453 | Unknown |
|  | SNOG_11397 | 204 | 0.9892 | 2.299064 | 9.19E-34 | 520.782 | 2598.135 | DUF4360 |
|  | SNOG_02244 | 159 | 0.9961 | 1.477338 | 7.95E-24 | 524.339 | 1486.912 | Nuclear transport factor like 2 |
|  | SNOG_06690 | 122 | 0.8006 | 1.321033 | 1.48E-05 | 485.8987 | 1296.928 | Unknown |
|  | SNOG_11929 | 302 | 0.9096 | 1.127803 | 2.90E-13 | 564.8313 | 1263.839 | Carboxylic ester hydrolase |
|  | SNOG_05030 | 114 | 0.9998 | 1.979014 | 1.19E-33 | 258.9124 | 1042.567 | Unknown |
|  | SNOG_01033 | 297 | 0.9787 | 1.229176 | 1.01E-08 | 353.7862 | 854.3281 | Apple domain |
|  | SNOG_13099 | 269 | 0.8591 | 1.024306 | 6.50E-06 | 343.9231 | 720.0231 | GH12 hydrolase |
|  | SNOG_00069 | 154 | 0.8132 | 1.016359 | 5.85E-10 | 340.7407 | 706.0248 | Necrosis inducing protein Nis1 |
|  | SNOG_15324 | 104 | 1 | 2.402469 | 6.51E-20 | 130.8397 | 704.7976 | Unknown |
|  | SNOG_00110 | 301 | 0.9997 | 3.495575 | 9.82E-61 | 61.74363 | 699.7316 | Unknown |
|  | SNOG_14877 | 208 | 0.9963 | 1.239108 | 1.67E-05 | 226.0309 | 565.0988 | Unknown |
|  | SNOG_04821 | 133 | 0.9373 | 1.149232 | 9.72E-09 | 230.3071 | 518.1454 | RALPH-like |
|  | SNOG_30829 | 226 | 0.7335 | 1.938823 | 8.41E-24 | 93.47433 | 364.9224 | Copper acquisition factor BIM1 |
|  | SNOG_00616 | 313 | 0.9948 | 1.120121 | 0.00022400 | 148.6516 | 342.956 | Chlorophyllase |
|  | SNOG_30113 | 296 | 0.9729 | 1.361109 | 2.65E-11 | 127.2852 | 342.401 | GH16 hydrolase |
|  | SNOG_15608 | 247 | 0.6907 | 1.02751 | 8.59E-05 | 133.725 | 288.0973 | Cutinase |
|  | SNOG_14857 | 238 | 0.994 | 1.303725 | 5.89E-08 | 97.24263 | 245.9543 | DUF4360 |
|  | SNOG_01569 | 324 | 0.8629 | 1.512704 | 1.65E-06 | 80.29615 | 238.7532 | GH17, Ubiquitin 3 binding |
|  | SNOG_40120 | 117 | 0.9993 | 1.62476 | 2.16E-09 | 57.33046 | 180.5937 | Unknown |
|  | SNOG_11853 | 138 | 0.997 | 1.30328 | 1.13E-06 | 64.16194 | 166.0219 | Unknown |
|  | SNOG_05588 | 343 | 0.9696 | 1.047068 | 4.21E-05 | 71.81932 | 156.4778 | Unknown |
|  | SNOG_15460 | 199 | 0.8384 | 1.091885 | 0.01140495 | 63.53512 | 153.8905 | Unknown |
|  | SNOG_07825 | 211 | 0.9999 | 1.390009 | 7.68E-07 | 48.25632 | 135.5342 | Chitin binding |
|  | SNOG_05309 | 277 | 0.5397 | 1.069146 | 0.01910536 | 42.32436 | 101.5459 | Peptidase |
|  | SNOG_14588 | 173 | 0.7208 | 1.446314 | 2.58E-05 | 29.48192 | 88.73187 | Unknown |
|  | SNOG_05629 | 111 | 0.9007 | 2.880919 | 2.21E-12 | 11.34205 | 85.76006 | Unknown |
|  | SNOG_15988 | 292 | 0.8552 | 1.7764 | 2.06E-05 | 17.82468 | 70.13389 | Unknown |
|  | SNOG_00801 | 334 | 0.9718 | 1.066381 | 0.00512434 | 27.6378 | 68.42403 | DUF1996 |
|  | SNOG_43715 | 103 | 0.9998 | 2.091915 | 0.00069420 | 11.71252 | 62.38947 | Hydrophobin |
|  | SNOG_14808 | 174 | 0.8636 | 1.453834 | 0.00030468 | 18.82044 | 60.0353 | Unknown |
|  | SNOG_13837 | 187 | 0.9601 | 1.040032 | 0.03770495 | 22.40941 | 57.62651 | DUF4360 |
|  | SNOG_11295 | 97 | 0.9999 | 1.994543 | 0.00035610 | 7.999843 | 37.82502 | Unknown |
|  | SNOG_30026 | 148 | 0.9984 | 1.531784 | 0.02658918 | 7.291745 | 27.16252 | Unknown |
|  | SNOG_44732 | 84 | 0.9998 | 1.917033 | 0.00754560 | 4.734496 | 23.16836 | Unknown |
|  | SNOG_01173 | 223 | 0.9967 | 4.62807 | 0.00156352 | 0.319379 | 14.30148 | Peptidase |
|  | SNOG_05548 | 340 | 0.9703 | 1.953009 | 0.02878011 | 1.473675 | 10.41415 | Kelch-like |
|  | SNOG_02352 | 114 | 0.8596 | 6.468327 | 0.00108152 | 0 | 7.656646 | Unknown |

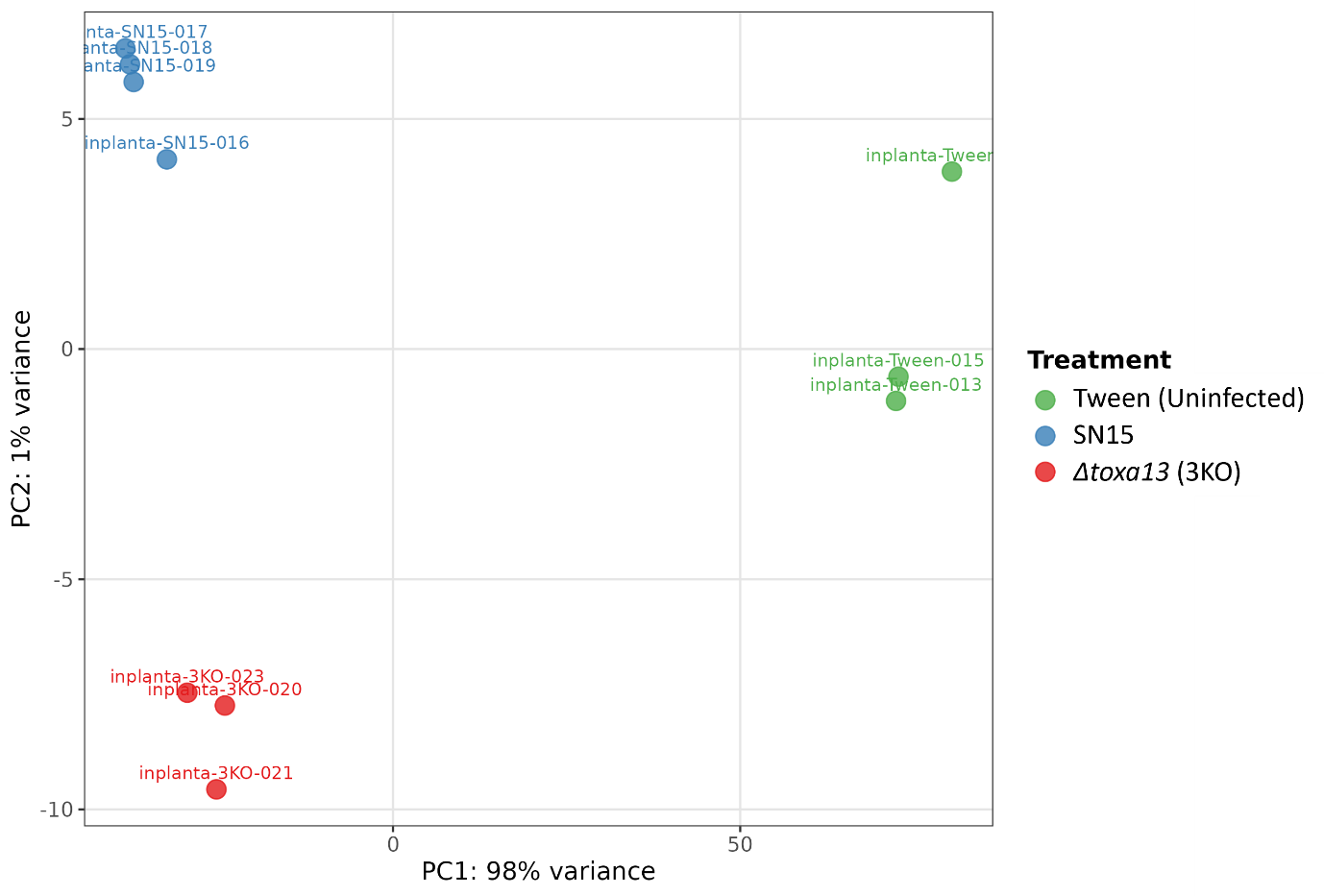

**Supplemental Figure S1:** Principal component analysis of transcriptomes of wheat-host against SN15 and *Δtoxa13* infection three-days post-inoculation, and uninfected/tween-treated leaves. Four biological replicates were used for SN15-infected leaves (Jones et al. 2019). Three biological replicates were used for uninfected/tween-treated leaves and *Δtoxa13*-infected (this study). All raw RNA sequencing reads generated in this study can be found in Bioproject PRJNA1479129.

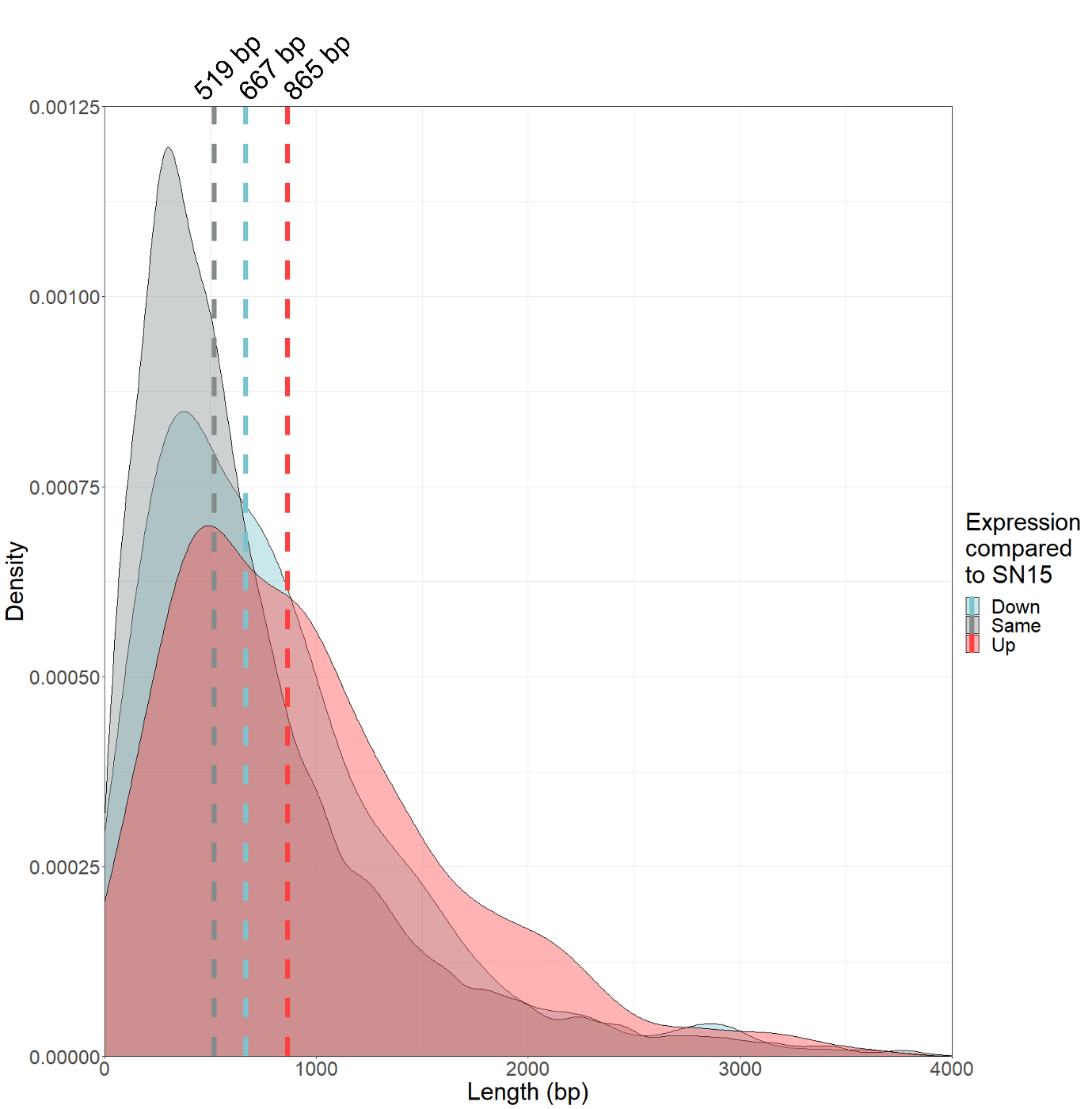

**Supplemental Figure S2:** Correlations in the upstream intergenic region (UIR) of all differentially regulated genes in *Δtoxa13 in-planta*. Distribution density plot of UIR lengths between down-regulated (Down), up-regulated (Up) and non-differentially regulated (Same) genes in *Δtoxa13 in-planta*, with the median lengths marked with a vertical dotted line. The median lengths significantly differ between groups according to the Kruskal-Wallis test and the *post-hoc* Dunn’s test with the Holm adjustment (adj. *p <* 0.001).

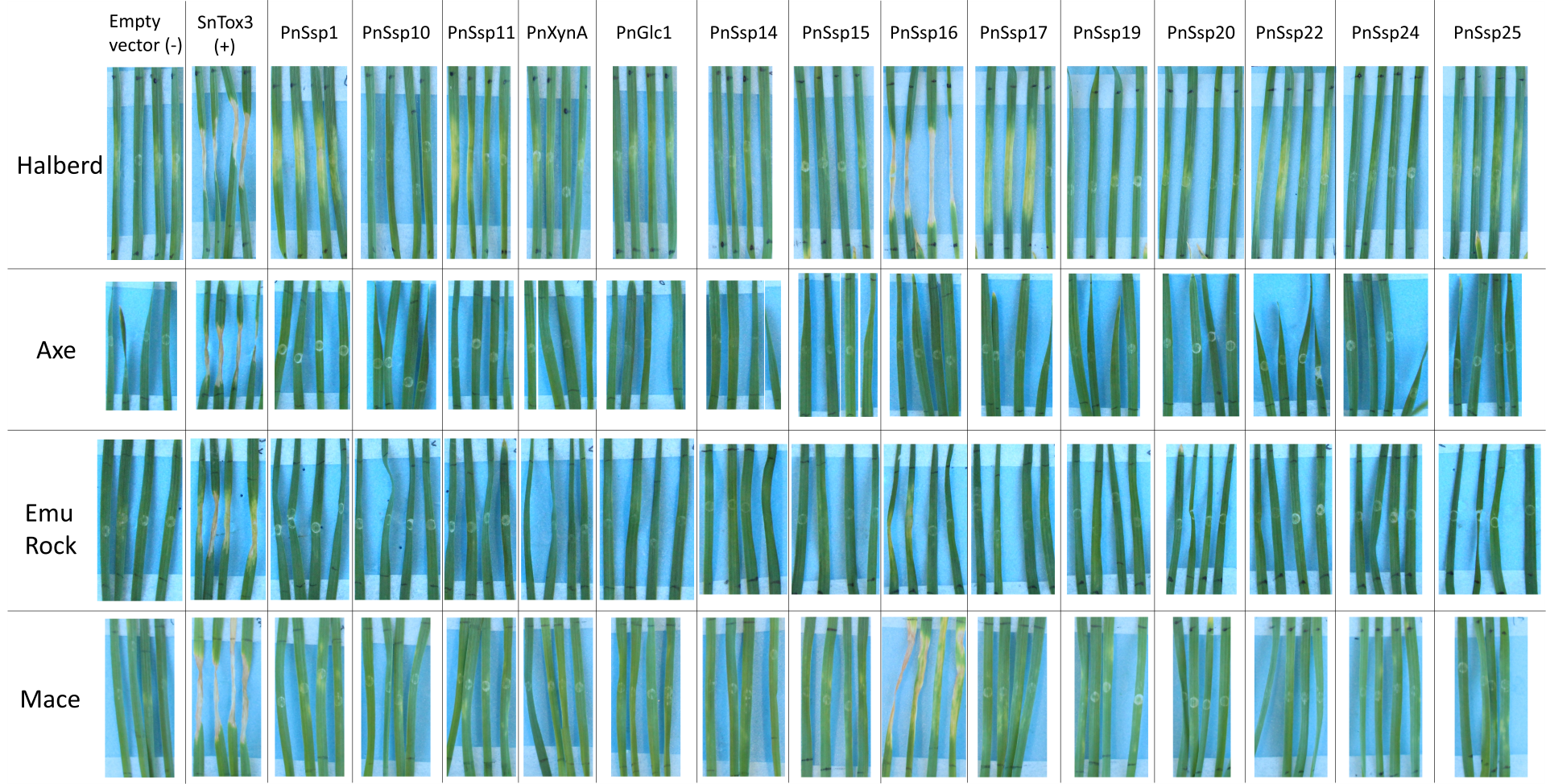

**Supplemental Figure S3:** *Pichia pastoris* culture filtrate infiltration of up-regulated candidate effectors from *P. nodorum Δtoxa13* into commercial wheat cv. Halberd, Axe, Emu Rock, Mace; seven days post infiltration.

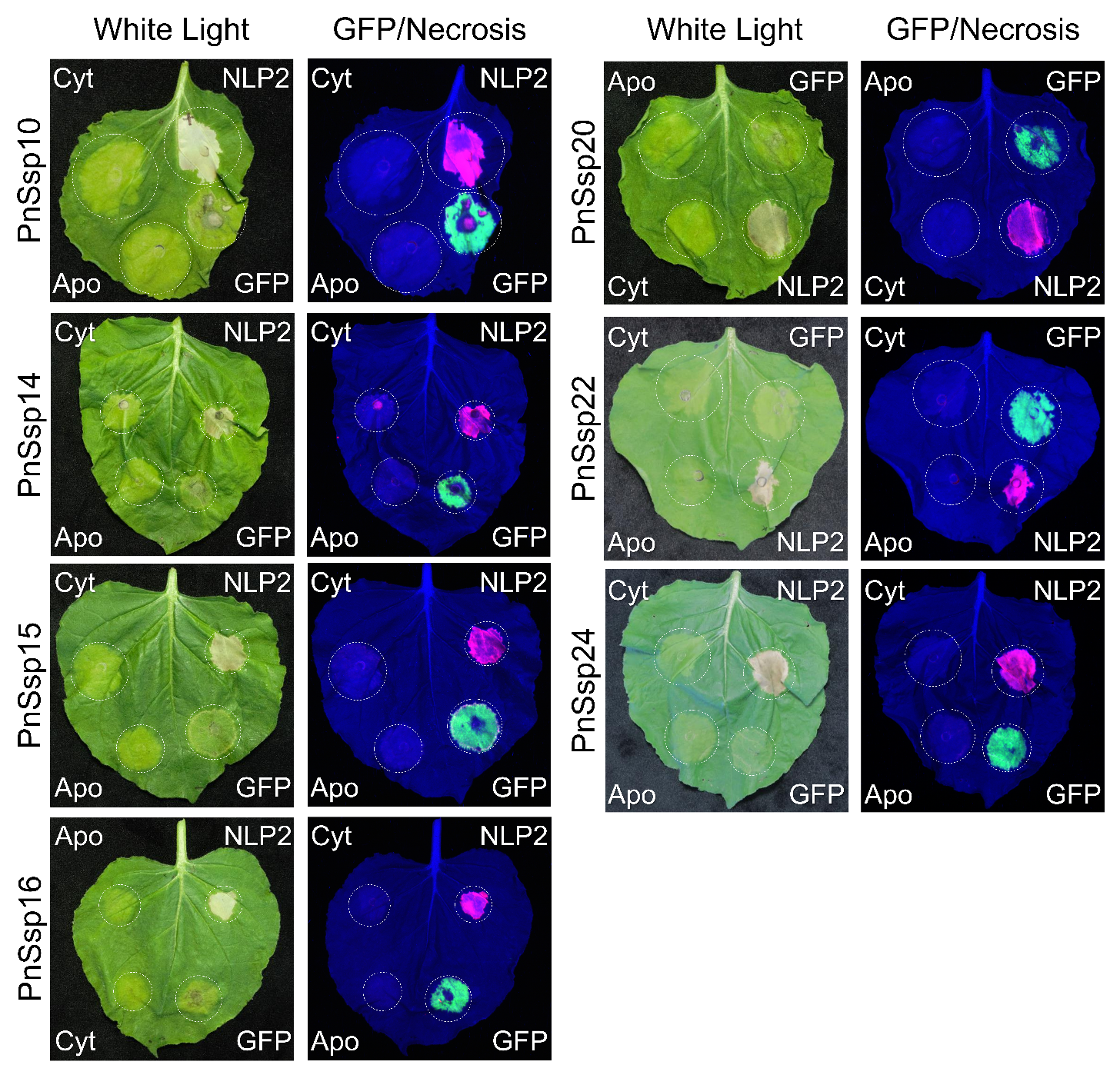
**Supplemental Figure S4:** Agroinfiltration of negative/non-host necrosis inducing up-regulated candidate effectors from *P. nodorum Δtoxa13* and expressed in SN15. Necrosis inducing effector candidates are shown in-text at **Figure 4**. “White light” shows representative *Nicotiana benthamiana* leaves agroinfiltrated with one of 14 candidate effectors and photographed 10 days post-infiltration. NLP2 (necrosis- and ethylene-inducing peptide 1 (NEP1)-like protein 2) and GFP (green fluorescent protein) are the positive and negative controls, respectively. Black names show no necrosis-inducing activity. The location of each agroinfiltration has been partially randomised. “Necrosis/GFP” shows the same leaf highlighting necrosis (pink) and GFP fluorescence (green). “Cyt” and “Apo” indicate cytoplasmic and apoplastic localisation of the candidate effectors, respectively.

**Supplemental Table S4:** Recombinant expression of SnTox267 infiltration into 78 different varieties of Australian commercial wheat cultivars. A score of 0 indicates insensitivity (white, no reaction); 1, slight chlorosis (green); 2, moderate chlorosis/slight necrosis (yellow); 3, moderate necrosis (orange); 4, extensive necrosis (red). (*one replicate of Baxter unsuccessful).

| **Cultivar** | **SnTox267 sensitivity score** | | |
| --- | --- | --- | --- |
| Axe | **3** | **3** | **4** |
| Baxter* |  | **0** | **2** |
| Beckom | **2** | **3** | **3** |
| Bolac | **2** | **2** | **3** |
| Borlaug 100 | **1** | **1** | **2** |
| Bremer | **1** | **1** | **2** |
| Brennan | **0** | **1** | **1** |
| Buchanan | **2** | **2** | **3** |
| Calingiri | **3** | **3** | **4** |
| Chara | **3** | **3** | **4** |
| Chief CL Plus | **4** | **4** | **4** |
| Cobalt | **2** | **2** | **3** |
| Coolah | **2** | **3** | **3** |
| Corack | **4** | **4** | **4** |
| Cutlass | **2** | **2** | **2** |
| Derrimut | **3** | **3** | **3** |
| DS Bennett | **0** | **0** | **0** |
| DS Darwin | **1** | **2** | **2** |
| DS Faraday | **2** | **2** | **3** |
| DS Pascal | **0** | **0** | **1** |
| EGA Gregory | **3** | **4** | **4** |
| EGA Wedgetail | **1** | **1** | **2** |
| Einstein | **3** | **4** | **4** |
| Elmore CL Plus | **3** | **3** | **4** |
| Emu Rock | **0** | **0** | **0** |
| Estoc | **1** | **1** | **3** |
| **Cultivar** | **SnTox267 sensitivity score** | | |
| Forrest | **3** | **4** | **4** |
| Grenade CL Plus | **1** | **2** | **2** |
| Harper | **1** | **3** | **3** |
| Hartog | **1** | **1** | **2** |
| Impress CL Plus | **0** | **0** | **0** |
| Kord CL Plus | **1** | **2** | **2** |
| Livingston | **3** | **3** | **3** |
| Longsword | **4** | **4** | **4** |
| LRPB Arrow | **4** | **4** | **4** |
| LRPB Beaufort | **0** | **0** | **1** |
| LRPB Cobra | **1** | **2** | **2** |
| LRPB Crusader | **1** | **1** | **1** |
| LRPB Dart | **2** | **2** | **3** |
| LRPB Flanker | **3** | **3** | **3** |
| LRPB Gauntlet | **3** | **3** | **3** |
| LRPB Havoc | **3** | **3** | **3** |
| LRPB Impala | **1** | **2** | **2** |
| LRPB Kittyhawk | **3** | **3** | **3** |
| LRPB Lancer | **2** | **2** | **3** |
| LRPB Lincoln | **4** | **4** | **4** |
| LRPB Mustang | **3** | **3** | **4** |
| LRPB Phantom | **2** | **2** | **3** |
| LRPB Reliant | **1** | **1** | **1** |
| LRPB Scout | **2** | **2** | **2** |
| LRPB Spitfire | **2** | **2** | **3** |
| **Cultivar** | **SnTox267 sensitivity score** | | |
| LRPB Trojan | **2** | **2** | **3** |
| Mace | **1** | **1** | **1** |
| Magenta | **3** | **3** | **4** |
| Naparoo | **0** | **0** | **0** |
| Razor CL Plus | **1** | **1** | **2** |
| RGT Accroc | **2** | **2** | **2** |
| RGT Calabro | **1** | **1** | **2** |
| RGT Zanzibar | **0** | **0** | **0** |
| Scepter | **1** | **2** | **2** |
| SEA Condamine | **3** | **4** | **4** |
| SF Adagio | **3** | **3** | **4** |
| Shield | **3** | **3** | **3** |
| SQP Revenue | **0** | **0** | **0** |
| Sunlamb | **4** | **4** | **4** |
| Sunmax | **2** | **2** | **2** |
| Suntime | **1** | **1** | **2** |
| Suntop | **2** | **3** | **3** |
| Sunvale | **3** | **4** | **4** |
| Sunzell | **1** | **2** | **2** |
| Tenfour | **0** | **3** | **3** |
| Tungsten | **1** | **2** | **2** |
| Wallup | **0** | **0** | **4** |
| Westonia | **1** | **1** | **1** |
| Wyalkatchem | **4** | **4** | **4** |
| Yenda | **2** | **2** | **3** |
| Yitpi | **1** | **1** | **1** |

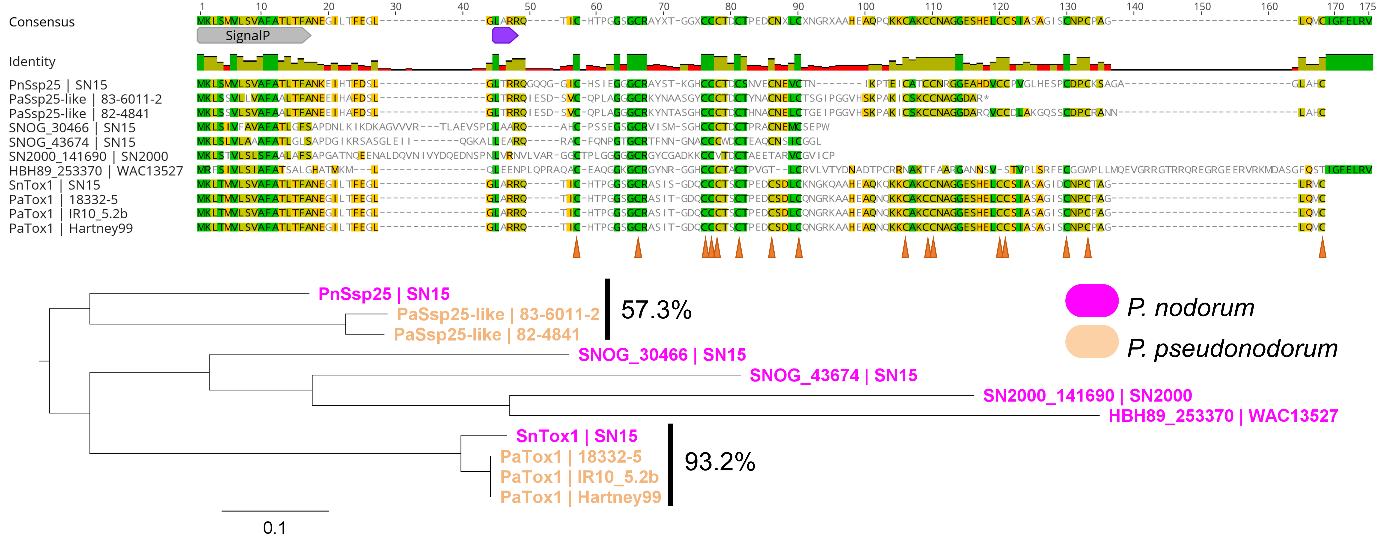

**Supplemental Figure S5:** Sequence analysis of candidate effectors PnSsp25. A protein sequence alignment of PnSsp25/SnTox1 homologs in *P. nodorum* and *P*. *pseudonodorum.* “SignalP” is the predicted signal peptide. The purple bar represents the location of the putative Kex2 cleavage site based on SnTox1/PnSsp25, and orange arrows indicate the location of 16 conserved cysteine residues in SnTox1 and PnSsp25. A phylogenetic tree showing the relatedness of PnSsp25/SnTox1 homologs in *P. nodorum* and *P*. *pseudonodorum*. The percentages next to the clades show the percentage sequence identity within the clades.

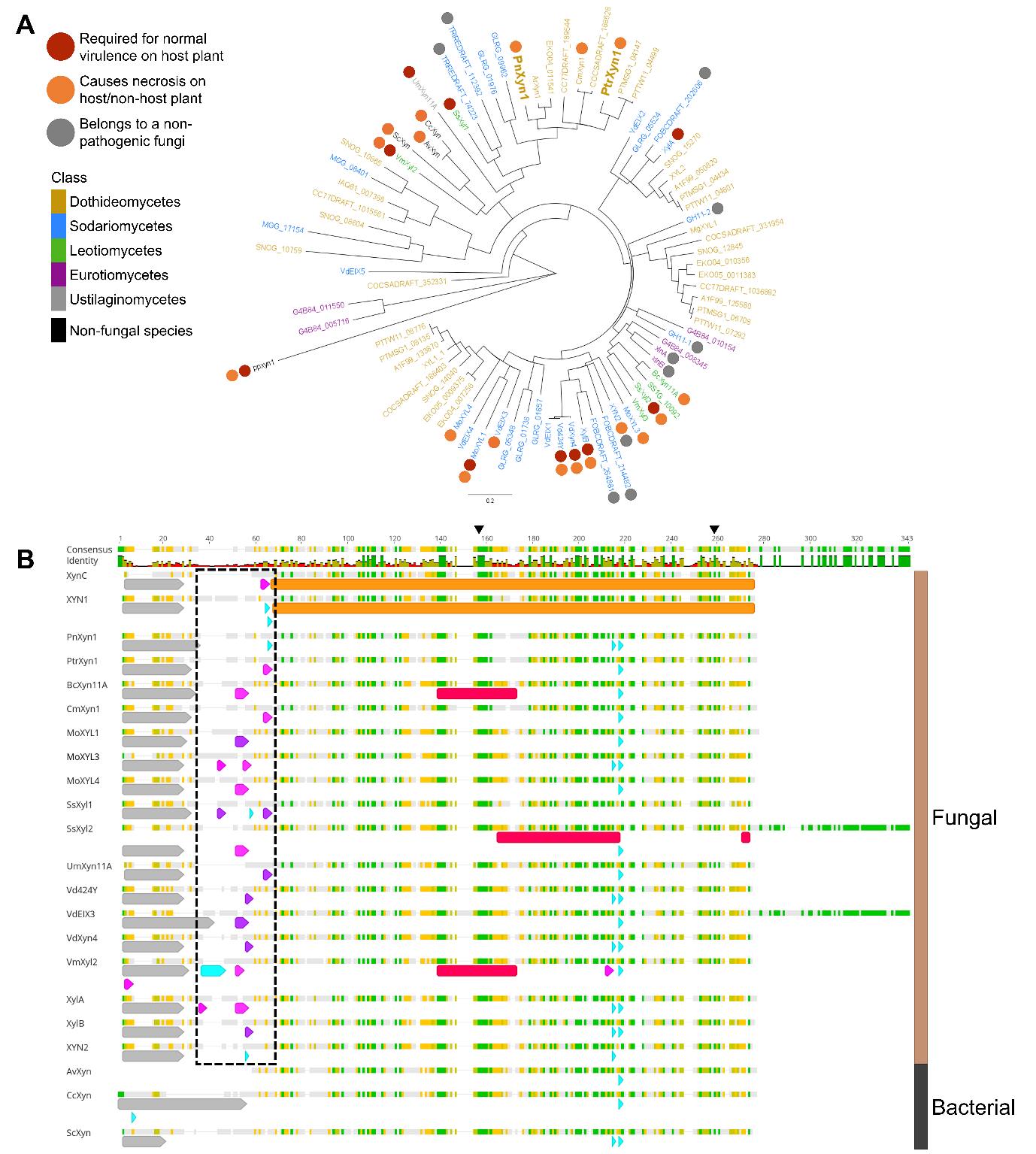

**Supplemental Figure S6:** Phylogenetic analyses and characterisation of the GH11 xylanase PnXyn1. (**A**) A phylogenetic tree showing GH11 xylanases in 23 fungal species, three non-fungal species and a necrosis-inducing GH10 xylanase (ppxyn1) as an outgroup. The nodes are coloured according to fungal classes. Red circles next to the nodes indicate xylanases required for full virulence by the fungi of origin. Orange circles indicate a necrosis-inducing activity on the host and/or non-host plant. PnXyn1 and the closely related PtrXyn1 are bolded. Grey circles indicate that the fungi of origin are non-pathogens. The scale bar indicates amino acid changes per site. (**B**) Sequence alignment of 22 GH11 xylanases, including structurally determined GH11 xylanases XynC (PDB: 1BK1) and XYN1 (PDB: 1XYN), showing putative N-terminal Kex2 cleavage sites only conserved in fungi. Grey represents predicted signal peptides. Putative Kex2 cleavage sites are coloured according to their recognition site – blue ((K/R)R), pink (LXXR) and purple (LX(K/R)R). The size differences of the putative Kex2 cleavage sites are due to gaps in the sequence alignment within the sites. The dotted box represents conservation of putative Kex2 cleavage sites in the N-termini of fungal GH11 xylanases. Red represents regions required or sufficient for necrosis-inducing activity on tested *Nicotiana* spp. The orange bars represent structurally determined regions of XynC and XYN1. The numbers show the length of the protein alignment. The brown and black bars on the right encompass GH11 xylanases of fungal and bacterial origin, respectively. The black triangles at the top of the alignment show the location of the conserved catalytic glutamic acid residues.
